# Hierarchical Community Structure of the Adult *Drosophila* Connectome Reveals Conserved Circuit Archetypes

**DOI:** 10.64898/2025.12.10.689094

**Authors:** Richard Betzel, Osmar Del Rio, Nathan Labora, Sophie Dvali, Bart Larsen, Christopher W. Lynn, Bratislav Misic, Linden Parkes, Olaf Sporns, Brenden Tervo-Clemmens, Maria Grazia Puxeddu, Caio Seguin

## Abstract

The community structure of connectomes supports functional specialization, adaptability, and cost-efficient wiring. However, little is known about communities in connectomes mapped at the level of individual neurons and synapses. Here, we analyze a whole-brain Drosophila adult connectome using a nested stochastic blockmodel to uncover its hierarchical community structure. Most of the roughly 1500 fine-scale communities–the smallest, and best resolved level of the hierarchy– are spatially compact, mostly assortative, and aligned with biological features. Nonetheless, we find evidence of nonassortative communities, spatially co-localized within the optic lobe and vision-processing pathways. Seeking “functional primitives”–small circuits with functionally narrow feature profiles–we use data-driven clustering to group communities into 45 archetypical meta-clusters based on their spatial, functional, and molecular properties, revealing modular building blocks from which larger, functionally diverse communities are composed. This work advances our understanding of how structure and function are organized in the fruit fly brain and highlights the value of statistical network models in interpreting nanoscale connectomes.

## INTRODUCTION

Connectomes are comprehensive network maps of the physical connections interlinking neural elements [1]. Understanding the principles by which these networks are organized and how their structure supports brain function and behavior is a central goal of neuroscience [2–5].

Connectomes can be modeled as graphs, wherein neural elements–neurons, neuronal populations, brain regions–are represented as nodes, and their pairwise interactions–synapses, projections, fiber tracts–as edges [4, 6, 7]. This abstraction enables quantitative analysis of brain network architecture, albeit at the cost of discarding some biological detail [8, 9]. Nevertheless, this network-centric framework has proven useful for identifying organizing principles of nervous systems across species and scales [8, 10–12]

One such principle is so-called “community structure”–divisions of a connectome into neuro-scientifically meaningful sub-networks, referred to as “modules” or “communities” [13– Communities are thought to support specialized information processing [17], enhance adaptability and evolvability [18, 19], and promote cost-efficient spatial embedding [20]. Indeed, modular organization appears to be a conserved feature of connectomes and can be observed across scales, ranging from the nervous system of *C. elegans* all the way to primates [21–28]

However, empirical studies of connectome community structure face two notable limitations. First, the majority of studies have been carried out on coarse-grained, mesoscale connectomes (projectomes), whose nodes correspond to entire brain areas [24, 29–32] This focus reflects the historical scarcity of neuron-resolved, whole-brain connectomes–*C. elegans* being a notable exception [33, 34]. As a result, it remains unclear whether the modular organization observed at mesoscale persists, transforms, or dissolves at finer resolutions (although, again, recent work has helped clarify this [22, 23]).

Second, most studies have focused on “assortative’– internally dense and externally sparse–communities. In theory, assortativity is an attractive property. In simulations, assortative architecture supports specialization of function [35], which is well-aligned with empirical observations in large scale functional networks showing that assortative communities circumscribe known functional systems [36]. Additionally, assortative structure promotes evolvability and robustness to perturbations [37], while facilitating cost-efficient spatial embedding [20]. The emphasis on assortative communities in neuroscience is reinforced by the widespread adoption of community detection heuristics like modularity maximization [38], which are incapable of detecting nonassortative communities [14]. On the other hand, purely assortative organization may be suboptimal, leading to approximately disjoint “islands” of neurons and limiting the cross-community integration of information flow. While previous studies of community structure in mesoscale [39–41] and small nanoscale connectomes [23, 26, 28] reported nonassortative communities, albeit at a low baseline rate, it is unknown whether large, whole-brain connectomes reconstructed at the nanoscale are organized according to assortative design principles.

Recent advances in dense electron microscopy, segmentation, proofreading, and semi-automated annotation have enabled the reconstruction of whole-brain, synapse-resolution connectomes in multiple species. These include the adult and larval *Drosophila melanogaster* [42–44], as well as partial reconstructions of zebrafish [25], mouse [45], and human brains [46]. These datasets present a unique opportunity to study the structural organization of neural systems at the scale of individual neurons and synapses [9].

In this study, we focus on the FlyWire reconstruction of a female adult *Drosophila* connectome [43]. This rich dataset consists of approximately 140,000 neurons and weighted, directed (chemical) synaptic connections. In addition to connectivity, the dataset is heavily annotated and includes information about neurotransmitter type, neuronal morphology, and expert-curated labels of anatomy, function, and cell type. Our goal is to leverage tools from network neuroscience to contribute complementary insights into the structural organization of the *Drosophila* nervous system, focusing specifically on its community structure.

Importantly, community structure and annotations potentially represent opposed perspectives on *Drosophila* organization. One emphasizes data-driven inference of neuronal groups based on their connection patterns, while the other emphasizes theory, biology, and domain knowledge. Our work seeks to explicitly compare these two viewpoints, highlighting points of consensus – communities that group together annotations with shared functional roles – along with disagreement – communities whose composition may be unanticipated given the extant literature and could suggest at novel circuits with unknown functions. To address these questions, we used the nested stochastic block model (SBM) [47] to estimate hierarchical communities. Unlike other algorithms popular in network neuroscience, SBMs are capable of detecting both assortative and nonassortative configurations [41]. We apply the SBM not merely to identify community boundaries, but to characterize the internal and external structure of those communities in detail. To this end, we quantify the assortativity of each community– i.e. whether its pattern of connectivity is primarily segregative, core-periphery, or disassortative. We find that most communities are, indeed, assortative, but a subset—particularly involving neurons linked to vision processing exhibits pronounced nonassortative structure. To interpret these communities, we conduct a series of enrichment analyses, testing how well aligned communities are with known cell types and functions. We observe a strong correspondence between expert annotations and data-driven community assignments, suggesting that a defining feature of many neurobiologically relevant annotations is their inter-connectivity. Finally, we integrate connectivity patterns, annotations, and spatial morphological information to construct a taxonomy of community “archetypes.” These meta-classes capture recurring motifs across the brain, offering a high-level view of how the *Drosophila* brain’s microstructure organizes into functionally and structurally meaningful units.

## RESULTS

We analyzed the connectome of a single adult female *Drosophila melanogaster*, publicly available through FlyWire (https://codex.flywire.ai/), using the v783 release [43]. This connectome contains *N* = 138, 639 neurons (more than 450 distinct cell types) and *M* = 15, 091, 983 edges, with a total of *M*_*w*_ = 54, 492, 922 synapses.

To detect communities within the connectome, we used a weighted and hierarchical stochastic block model (SBM) [47– a method well-suited for uncovering modular structures in large-scale networks. The SBM approach is particularly effective for identifying nested hierarchical organization within the brain, a concept aligned with the long-standing belief that many natural and engineered systems are hierarchically modular [50]. This type of hierarchical connectivity has been observed in other species, such as primates and rodents, and it suggests a general principle of network organization that extends across different species and scales of biological complexity [13, 23]. In the following sections, we describe the communities in detail, link them to neuronal annotations, and posit that, based on their architectural features, they can be grouped into functionally narrow archetypes at the finest hierarchical scale and combined into broader, distributed communities at coarser levels of the hierarchy (see Fig. 1 for a broad overview of our approach).

**Figure 1.**
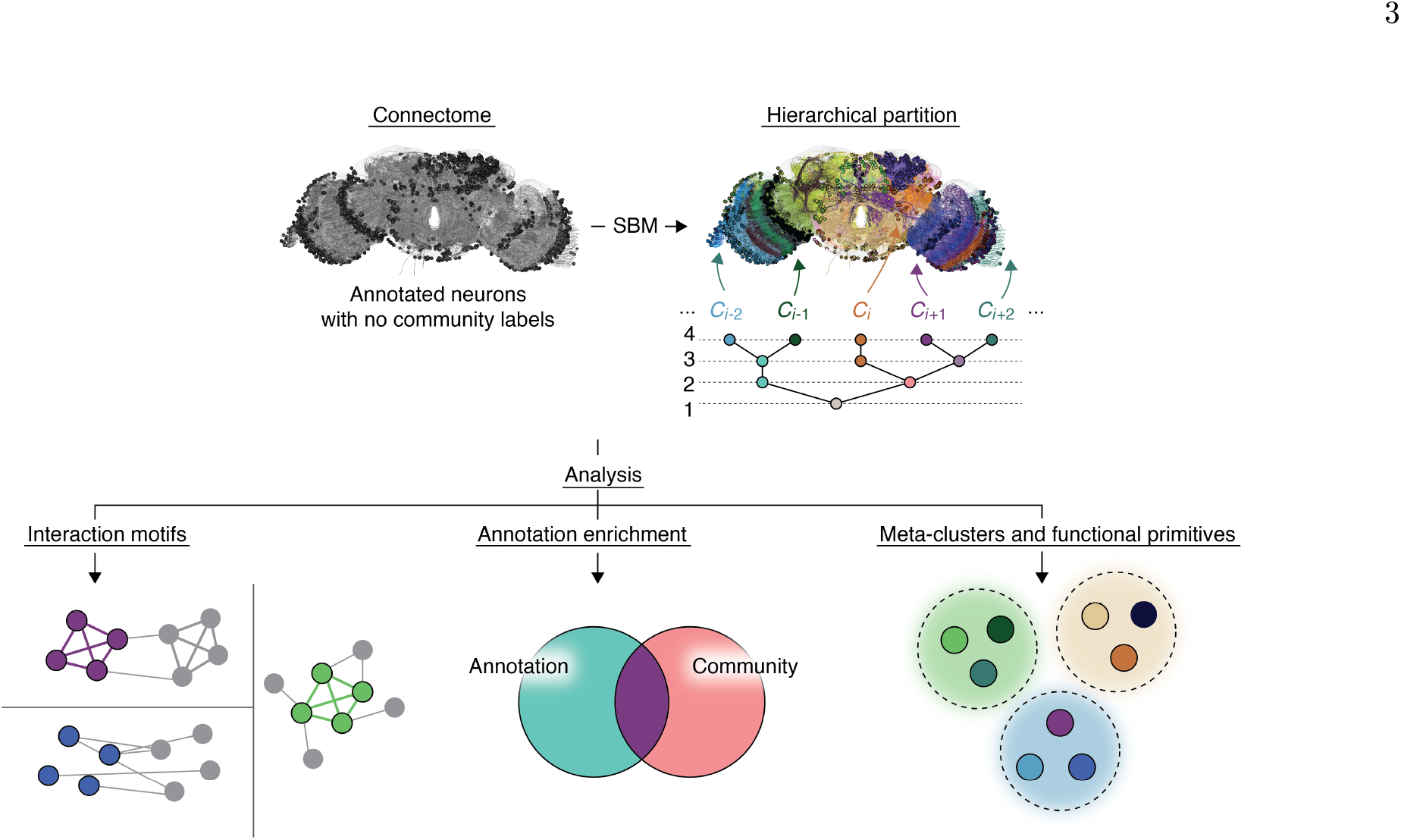
Analysis schematic. We study neuron-to-neuron connectivity from a female, adult *Drosophila melanogaster*. We use a blockmodel to partition neurons into non-overlapping, hierarchically organized communities. Our primary aim is to better understand the character of these communities. We do so using three distinct analytic approaches that address outstanding questions in network and computational neuroscience: Are communities in fully developed connectomes at the nanoscale strictly assortative? Do communities overlap with expert annotations of neurons? Can communities be classified into distinct archetypes based on their local features?

### Community Statistics

The SBM partitioned the *Drosophila* connectome into multi-level, nested communities. Here, we describe that partition and those communities in terms of low-level statistics. The consensus partition estimated using the SBM divided the connectome into 11 hierarchical levels. The number of communities at each level was: 1449, 498, 207, 114, 60, 35, 18, 10, 5, 3, and 2. However, upon visual inspection, we noted that the coarsest partitions (with only 2 and 3 communities, respectively) grouped the right optic lobe (along with visual sensory neurons) and central brain into the same community, while left-lateralized visual structures formed a separate community. Although asymmetric anatomy and function in *Drosophila* have been previously described, its nervous system is generally bilaterally symmetric [51, 52], and this bilateral symmetry is expected to be preserved at the resolution of communities. Therefore, we discarded the top two levels of the hierarchy, as they lacked neurobiological realism in this respect. In contrast, the five-community partition successfully separated the central brain from both left and right optic lobes. We therefore treated it as the highest valid level of the hierarchy.

To visualize community structure, we color-coded neurons by their community assignments and presented those alongside the weighted and directed connectivity matrix, ordered by community label (Fig.2A,B). We also show select communities in anatomical space, serving to highlight the spatial compactness and co-localization of neurons within the same community (Fig.2C). In this purely illustrative example, we chose communities that align with `super_class` annotations, which serve as a point of reference for the functional relevance of the detected modules. We note also that most communities exhibited a high level of bilateral symmetry–i.e. they spanned both the left and right sides of the brain or, if they were lateralized (as in the case of many optic lobe communities), there were approximate homotopic matches based on visual inspection.

Although sparse, the complete connectome contains roughly 140,000 neurons, hindering visualization and interpretation. To improve both, we generated a coarse-grained version of the connectome, where synaptic connections were aggregated by community. This network consisted of 1449 nodes (each corresponding to a fine-scale community). This renormalization and coarse-graining also facilitates the shifting of focus from individual neurons to the higher-level organization of communities. For instance, using this module-level representation of the connectome, we calculated a community-level coassignment matrix, whose entries represented how frequently (a count) fine-scale communities were grouped together at coarser hierarchical levels (Fig. 2D). This matrix provides insight into the modular relationships between communities at different scales and underscores the hierarchical structure of the connectome.

**Figure 2.**
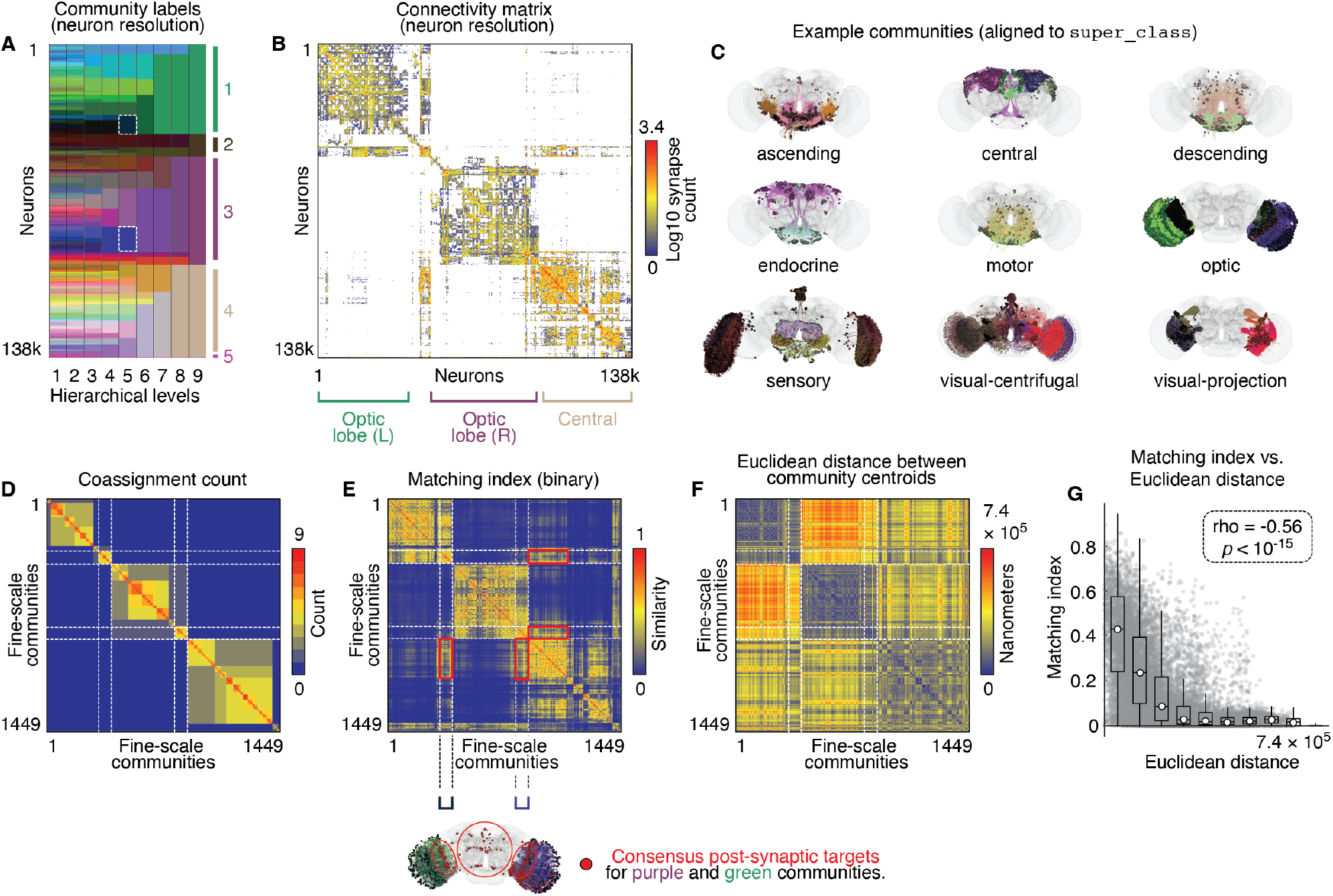
Community description. The SBM partitioned neurons into a hierarchy of communities. (*A*) Each fine-scale community received a unique color. Larger communities were given the mean color of children communities. (*B*) Whole-brain connectivity matrix at neuron-resolution (rows/columns ordered by community). (*C*) Unions of small communities aligned with `super_class` annotations. We aggregate neurons by their fine-scale community assignment and show: (*D*) community coassignment matrix; (*E*) community “matching index” (similarity of incoming/outgoing connections). The two coarse-scale optic lobe communities exhibited low overlap with the central brain community. However, there a select set of central brain communities exhibited high levels of overlap with left/right optic lobes. In the margin (below panel *E*) we show those communities in red. (*F*) Euclidean distances between all pairs of community centers of mass. (*G*) Matching index decays with center of mass Euclidean distance (*ρ* = −0.66; *p <* 10^−15^).

Additionally, we calculated the “matching index” for all pairs of communities. This measure quantifies the overlap between two communities’ connectivity profiles (in this case, accounting for both incoming and outgoing connections) [53] (Fig. 2E). The matching index is generally considered a measure of functional similarity, in line with the hypothesis that similar connectivity implies similar function, and is bound to the unit interval [0, 1] [54, 55]. Indeed, we find that matching indices in the coarse-grained *Drosophila* connectome are related to broad anatomical labels. For instance, the left and right optic lobes (green and purple communities) and central brain (brown) all exhibit high levels of within-area matching, but relatively low matching between areas.

The observation that the left and right optic lobes have relatively low matching with one another suggests that spatial constraints may also be an important determinant of connectional similarity, as these regions are located on opposed sides of the *Drosophila* brain despite having the same functional role. To further investigate the effect of spatial embedding, we calculated centroids– centers of mass–for each community and computed their pairwise Euclidean distances (Fig. 2F). While connection weight (normalized synapse count) between communities was unrelated to distance (Pearson, Spearman correlation coefficients; *r* = 0.0065, *ρ* = 0.0224 for community pairs linked by at least one synapse), inter-community matching exhibited a strong anti-correlation (*r* = − 0.55, *ρ* = − 0.59 for all community pairs), suggesting that connectivity overlap decreases as communities grow further from one another in space, a feature that the coarse-grained *Drosophila* connectome shares with other meso-scale connectome reconstructions [56– 59]. We speculate that the absence of a strong anti-correlation between synapse count and distance could reflect wiring specificity at neuron-resolution that is decoupled from spatial proximity and not evident in mesoscale connectomes [60]. We note, however, that neurons features extensive arbourization far from their respective soma. All center of mass estimates were based only on soma locations and do not account for neurons’ morphology.

In the **Supplementary Materials**, we provide a more comprehensive summary of community statistics across hierarchical levels, including community size, within-community synapse density, spatial compactness, and the relationship between community density and size (Fig. S1). Additionally, we performed analogous analyses at the neuron level (as opposed to the level of communities). We found that matching index was still anti-correlated with inter-soma distance, albeit more weakly (*ρ* = − 0.35). Also at the neuronal level, we found that connection probability decays monotonically with inter-soma distance (Fig. S2), although, among neurons connected by at least one synapse, there was no strong relationship between inter-soma distance and the number of synapses by which they were connected (*r* = 0.02; *ρ* = 0.04). In summary, our results provide statistical evidence that the *Drosophila* connectome is organized into nested communities with distinct spatial and statistical profiles. Many of these features are directly analogous to those identified in other connectome datasets, including the non-human mammalian brains [61, 62] and the human brain [24].

### Community Segregation and Interaction Motifs

In the previous section, we presented high-level summary statistics of communities in the *Drosophila* connectome. In this section, we focus on one particular feature in greater detail, community assortativity. Assortativity refers to the propensity of nodes to preferentially connect to other nodes in the same community. Networks with high levels of assortativity exhibit high levels of community segregation and, when the rows and columns of their connectivity matrices are sorted by community label, display the classical “block diagonal” structure. This type of modular organization has long been hypothesized to support specialized information processing [35] while buffering perturbations [63] and promoting evolvability [19]. Indeed, there exists a large body of literature documenting assortative community structure in connectomes at all spatial scales [22, 24, 25, 29, 30, 58, 64–67], suggesting that brains may be optimizing their organization to obtain these benefits.

To date, however, most studies of connectome community structure have relied on community detection tools that are biased towards the discovery of assortative communities–e.g. optimizing the classical modularity heuristic [38]. It is therefore unclear whether reports of exclusively assortative connectomes reflects methodological limitations or true features of connectomes. Here, however, we obtain community estimates using a stochastic blockmodel. In principle, blockmodels can detect nonassortative communities should they exist and are therefore less likely to generate a biased estimate of assortativity.

Here, we assess the level of assortativity in the network by examining community interaction motifs [41, 68]. Specifically, for every community dyad composed of communities *r* and *s*, we characterized their interaction as *assortative* if neurons in both communities preferentially connected to neurons within the same community – i.e., if within-community synapse densities, *d*_*rr*_ and *d*_*ss*_ both exceeded the between-community density, *d*_*rs*_. On the other hand, a dyad was considered disassortative if *d*_*rs*_ *>* max[*d*_*rr*_, *d*_*ss*_] or *core-periphery* when *d*_*rr*_ *> d*_*rs*_ *> d*_*ss*_ (in this case *r* is the core and *s* is the periphery). Whereas assortative interactions reflect segregated, specialized processing motifs, these nonassortative interactions reflect integrative and directed flows of information, as in “sender-receiver” or “feed-forward” processing [20].

We applied this classification scheme to every pair of communities across all hierarchical levels. Our focus, however, is on the finest resolution in which neurons were partitioned into 1449 communities, resulting in a total of 1449 *×* 1448 = 2, 098, 152 community pairs (excluding self-interactions). In addition to the three interaction motifs described above, we also included an “ambiguous” classification, which occurred when density measures were equal. In general, ambiguous motifs were infrequent and generally involved very small communities, thereby having negligible impact on the overall results.

We found that ≈ 94% of community interactions were assortative (Fig. 3A,D), suggesting that at neuron resolution, the *Drosophila* nervous system is largely composed of segregated modules. However, as with previous applications of blockmodels to connectome datasets [23, 39, 41, 68], we also observed a non-negligible fraction of nonassortative interactions: fewer than 1% were classified as disassortative, but nearly 2% as core-periphery. Although these nonassortative motifs were relatively infrequent, they nonetheless exhibited a high degree of anatomical specificity, concentrating in the optic lobes and among visual centrifugal/projection neurons (Fig. 3H,I).

**Figure 3.**
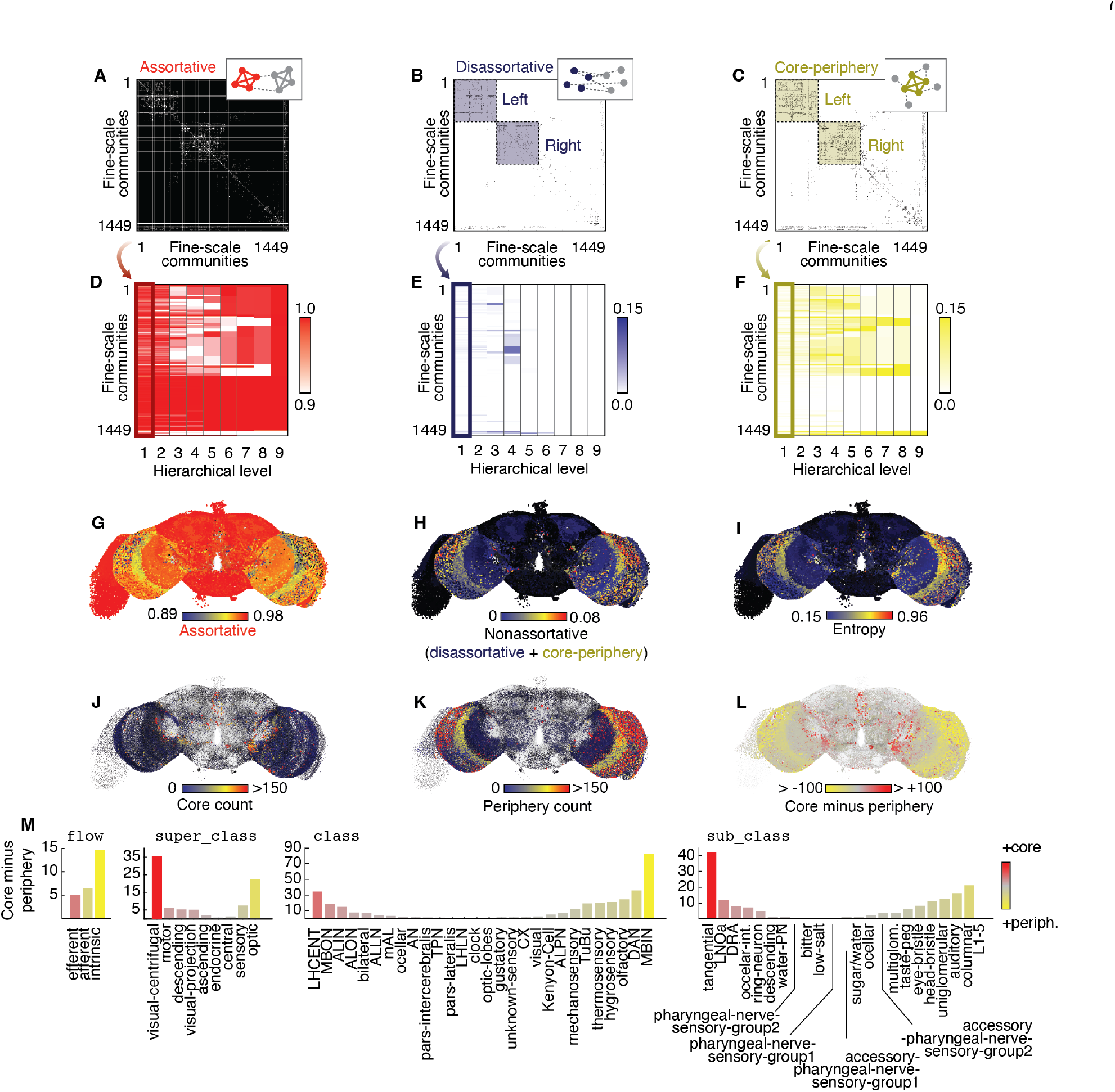
Community assortativity. We classified all pairs of fine-scale communities as *assortative, disassortative*, or *core-periphery*. Panels *A, B*, and *C* display categorizations at the finest resolution. Colored entries indicate pairs of communities assigned to the corresponding interaction motif. We highlight left/right optic lobes in the disassortative and core-periphery matrices, outlining the communities that comprise those structures. Panels *D*-*F* show interaction motif density across all hierarchical levels. Panels *G* and *H* show assortativity and non-assortativity in anatomical space. (*L*) We calculated an entropy measure for each community; greater levels of entropy indicated a greater diversity of interaction motifs. Panels *J* and *K* show core and periphery scores separated, respectively. We show the difference in core and periphery scores in (*L*), but also grouped based on annotation (*M*).

Broadly, this composition of neurons is involved in processing of visual information and relaying it to neurons in the central complex, where it is integrated and contributes to higher-order behavior, e.g. navigation and decision-making [69–71]. These results hint that non-assortativity may be a key architectural feature involved in the early stages of Drosophila sensory processing (vision, in this case), allowing for signals to move rapidly from peripheral sensors to core-like secondary targets before reaching more segregated post-secondary targets, where it is used to initiate and coordinate complex behavior[57, 58, 72].

### Contextualizing Communities with Annotations

To this point, we have calculated community statistics and shown results challenging the conventional view that neurons are organized into strictly assortative communities. However, one of the primary advantages of nanoscale connectomics is the depth of meta-data that accompanies connectivity information. Neurons are profiled along multiple dimensions, ranging from cell types to anatomical region to putative function. These annotations serve as indispensable guideposts, facilitating the contextualization and grounding of connectivity-based analyses in detailed neurobiology. Here, we leverage these rich annotational as means of validating and ascribing meaning to the detected communities.

First, we used annotations as part of a “community enrichment analysis.” Akin to functional enrichment analyses in genomics [73], which quantify the overlap of gene sets with biological functions, we assessed the extent to which fine-scale communities (1449 in total) overlapped with with cell type, anatomical, and functional annotations (Fig. 4A). Because raw overlap scores are biased by community and annotation size (all things equal, larger communities and annotations will tend to exhibit greater overlap), we expressed the enrichment scores as standardized measures, z-scoring them with respect to a null distribution generated by randomly permuting the order of annotations (1000 randomizations; Fig. 4B).

**Figure 4.**
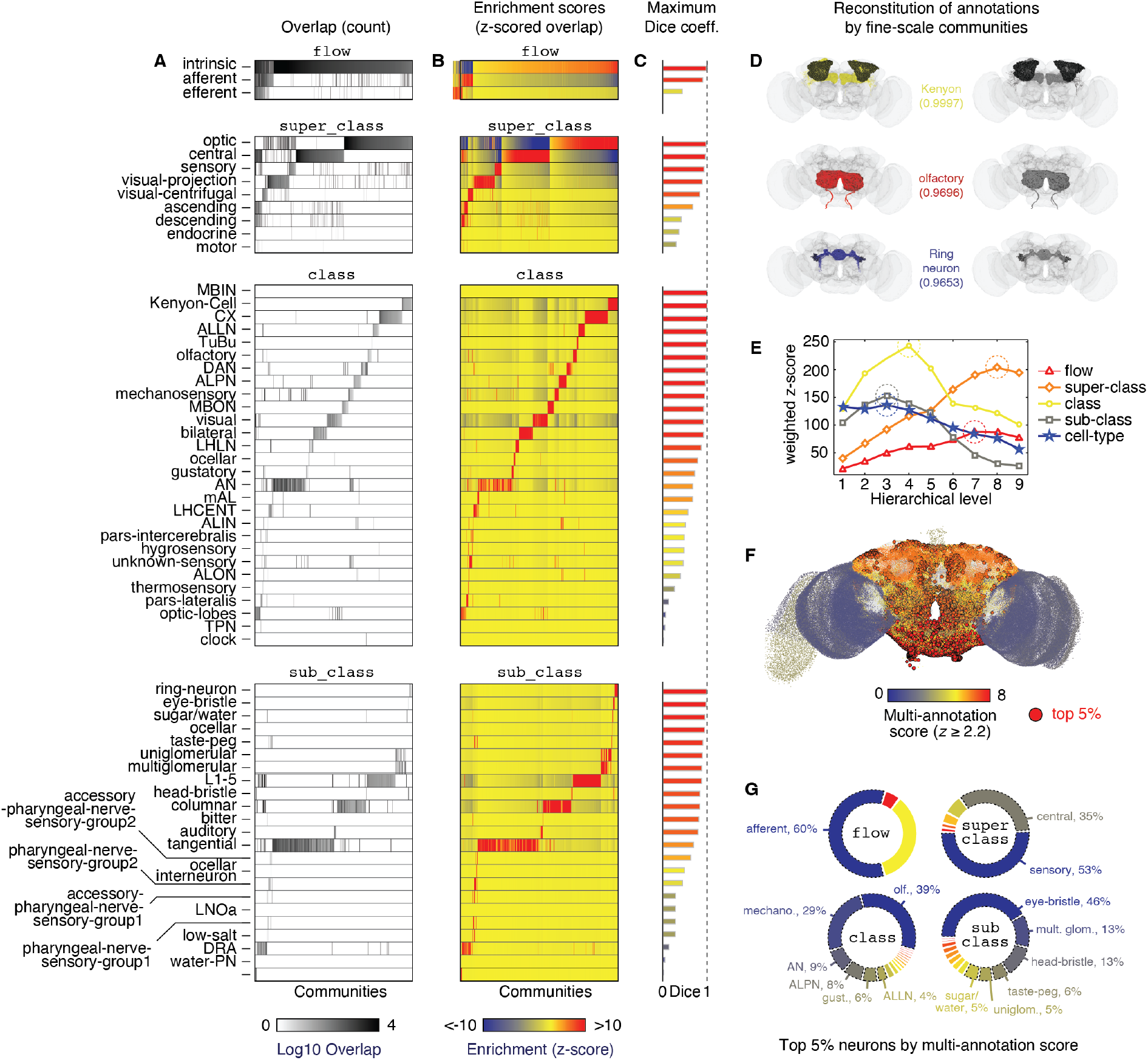
Community enrichment through annotations. We compared fine-scale communities against four levels of annotations. (*A*) Logarithm of overlap counts. Darker colors indicate greater levels of overlap. (*B*) To contextualize these measures, we compared and z-score the overlap counts against a permutation-based null model. Here, we show z-scores – enrichment scores – for each overlap measure. (*C*) We asked how well the union of subsets of fine-scale communities could approximate certain annotations (overlap measured as a Dice coefficient). We show the optimal Dice coefficient obtained for each of the annotations show in panels *A* and *B*. (*D*) Example annotations and best fit reconstructions from fine-scale communities. (*E*) Peak enrichment scores for different annotation categories. (*F*) Multi-annotation enrichment scores. The number of annotation categories for which different collections of neurons are enriched (*z >* 2.2). (*G*) Composition of top 5% neurons ordered by multi-annotation enrichment score across different annotations.

Interestingly, we found that the majority of communities exhibited non-trivial levels of overlap with annotations, suggesting that communities, which were derived from connectivity data alone, broadly recapitulate established neurobiological and functional categories. More interestingly, we found that the communities tended to exhibit preferences for singular annotations. As a demonstration, we measured the difference between a community’s dominant annotation (its top-ranked enrichment score) with its second-best score (Fig. S5). We then repeated this analysis, comparing secondary to tertiary enrichment scores, tertiary to quaternary, and so on. We found that the difference between dominant and secondary annotations was significantly greater, on average, then all other gaps (*t*-test; *p <* 10^−15^), an effect that held for different annotation types. These observations suggest that most communities were well aligned to a single annotation and largely homogeneous (97.2%, 98.2%, 96.9%, 96.4%, and 94.7% of all communities had enrichment gaps greater than two for `flow, super_class, class, sub_class`, and `cell_type` annotations). As a clear example, consider the `super_class` subplot in Fig. 4A. The communities that overlapped most with the “optic” label displayed almost no overlap with other annotations (in the overlap plot, their entries in other rows are close to zero and their enrichment scores in those rows are neutral or negative).

Nonetheless, not every community exhibited a clear annotational preference. Many exhibited multiple supra-threshold enrichment scores (here, we adopt a threshold of *>* 1.96 or roughly a *p <* 0.05). These multi-annotation communities exhibited a high degree of spatial specificity and were composed largely of afferent, non-visual sensory neurons as well intrinsic neurons in the central brain (Fig. 4F,G). This particular composition – olfaction and mechanosensation paired with other other intrinsic neurons – is of functional relevance, as these same neurons form circuits that support multisensory integration–e.g. detection and reaction to mates, food, or predators as well as feeding, grooming, and escape [74, 75]. These findings suggest that, although most communities show a clear annotational preference, deviations from this principle maybe functionally relevant and useful signatures of established circuits.

We also performed two additional analyses. First, we wanted to assess, theoretically, whether it was even possible to recover annotations from communities. That is, could we generate a set of neurons, *C* = {C_*i*_ ∪ *C*_*j*_ ∪ … ∪ *C*_*k*_ ∪ *C*_*l*_}, based on their community assignments such that *C* maximally overlapped with a specific annotation? The results of this analysis would serve to highlight annotations that are well-captured by connectivity-driven communities, but also those that are poorly represented. To this end, we used a simulated annealing algorithm to aggregated communities in order to maximize the Dice coefficient with respect to each annotation (Fig. 4C). Dice coefficient measures the overlap between two sets; a value of 1 indicates perfect overlap, while a value of 0 indicates that the two sets are dijoint. Surprisingly, we were able to almost perfectly recapitulate certain annotations. For instance, the Dice coefficient between communities and Kenyon cells was 0.9997 (of the 5177 annotated Kenyon cells, the optimal combination of communities correctly included them all, but along with three non-Kenyon cells; Fig. 4D, top). We observed similar performance when recovering olfactory and ring neuron annotations (0.9696 and 0.9653, respectively; Fig. 4D, middle and bottom). More generally, when we repeated this analysis for all flow, `super_class, sub_class`, and `class` annotations and weighted Dice coefficients by annotation size, we obtained mean scores of 0.9505 ± 0.0017, 0.9114 ± 0.0005, 0.9275 ± 0.0009, and 0.8458 ± 0.0004, respectively (mean ± standard deviation over repeated runs of the optimization algorithm), suggesting that annotations and communities are closely related.

However, we also observed that some annotations were difficult to recover. For instance, the “efferent” annotation (under `flow`) achieved a more modest Dice coefficient of 0.44. Similarly, `super_class` annotations for “motor” (0.30), “endocrine” (0.37), and “descending” (0.38) neurons also performed less well. Notably, these annotations were among the smallest in terms of size, underscoring the fact that very small annotations, which are more common in `sub_class` and `class` categories, may be challenging to recover from communities that are, by definition, composed of relatively large groups of neurons.

Rather than optimizing overlap outright, we also asked whether overlap peaked at particular hierarchical levels. That is, is there a characteristic size of community that is best aligned with different annotations? Specifically, we calculated the weighted enrichment score for each annotation type as a function of hierarchical level. Interestingly, we found considerable variability in terms of the level at which enrichment scores peaks. The peaks for `flow, super_class, class, sub_class`, and `cell_type` were at levels 7, 8, 4, 3, and 3. These observations suggest that different annotations may be preferentially expressed at different levels of the hierarchy and, importantly, these preferred levels are not directly linked to the coarseness of the annotations, themselves–e.g. `flow` peaks earlier than `super_class`, despite having only three categories (compared to the nine `super_class` categories).

Note that, as an intermediate step in estimating a global (whole-brain) enrichment score for each of the four annotation types, we calculated annotation-specific maximum Dice coefficients across hierarchical levels. That is, for each annotation, we calculated its maximum overlap with respect to any of the communities at each level of the hierarchy (Fig. S3). We then compared these overlap scores to the optimized overlap in Fig. 4C. We found that a significant fraction of annotations were as well approximated by single communities at coarser levels of the hierarchy as they were by the optimization procedure (21 of the 65 annotations), and that more than half (34 of 65), were within 75% of their optimized coefficient. These observations emphasize the multi-scale nature of community structure and its ability to connect connectivity with annotations at different characteristic scales. However, because many annotations were *not* recovered by any individual community and fall short of their optimized overlap scores, these observations also highlight the misalignment of communities with annotations. The optimized scores were achieved only by aggregating communities from separate branches of the hierarchy.

We further explored the alignment of annotations with communities by calculating the frequency with which different annotations were coassigned as part of the same community. If communities were perfectly homogeneous in terms of the annotation composition, then the annotation *×* annotation coassignment matrix would only have entries along its diagonal – i.e. dissimilar annotations would never co-mingle within the same community. On the other hand, deviations from this identity matrix might suggest an alternative ontology of annotations driven by connectivity information. We performed this analysis, calculating how frequently pairs of annotations were coassigned to the same fine-scale community (Fig. 5). We found that annotations preferentially co-clustered with themselves; the on-diagonal entries were significantly greater than the off-diagonal entries for each coassignment matrix (*t*-test; *p <* 10^−15^). Nonetheless, we observed some interesting structure. For `class` and `sub_class` categories, the coassignment graphs fragmented into disconnected components. In the case of class annotations, the components coincided with different sensory modalities; one component was composed of mechanosensory annotations (e.g. eye and head bristle neurons), another composed of olfactory neurons (uni-/multi-glomerular cells), and yet another for neurons associated with vision processing (tangential, columnar, laminar, and ocellar neurons). We also observed analogous fractionations in the `sub_class` coassignment graph, wherein Kenyon, mushroom body input, and tubercle–bulb neurons were fully disconnected from the rest of the graph. Even the connected component encoded sensible relationships; bilateral, visual, optic lobe, and ocellar neurons formed a conspicuous chain that was sparsely integrated into the connected component graph. Similarly, hygrosensory, olfactory, and thermosensory neurons formed a small clique of mutually co-clustered sensory neurons.

**Figure 5.**
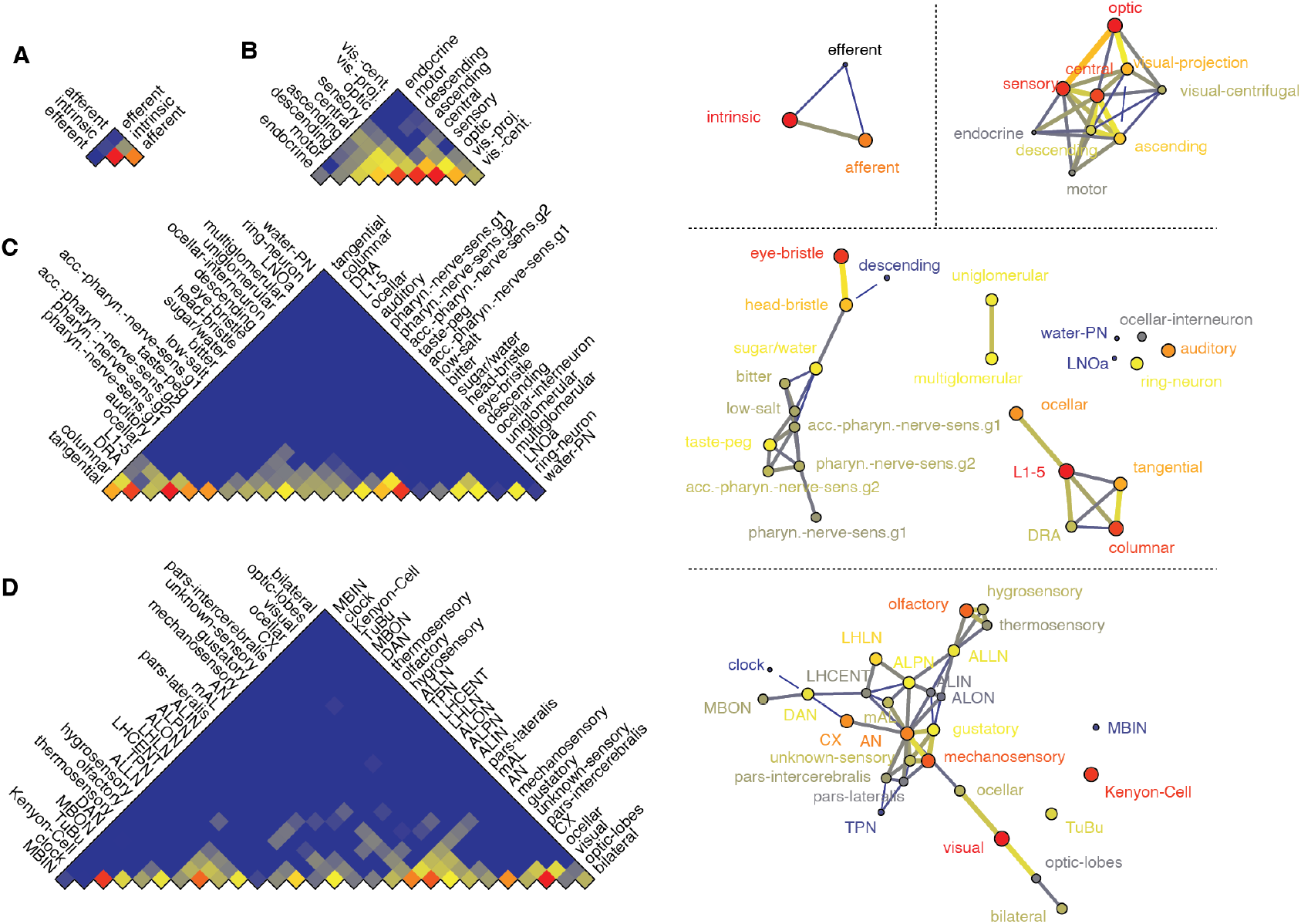
Neuronal coassignment by annotation. The SBM partitions neurons into communities. Earlier, we asked to what extent annotations are concentrated within communities. We can flip this and ask the complementary question: to what extent do neurons in the same community concentrate within any annotation? Here, we carry out this analysis from the perspective of four annotation levels: `flow, super_class, class`, and `sub_class`. The logarithm of coassignment counts are shown in panels *A*-*D*. From these, we can also depict these counts as graphs – annotations are linked to one another based on the frequency of their coassignments. We show these graphs using force-directed embedding in the right sub-panels.

Finally, as part of a supplementary analysis, we explored links between community structure and the `hemilineage` annotation, which labels neurons based on the neuroblasts from which they originated [76]. Rather than considering the whole brain, we focused specifically on communities that overlapped with Kenyon cells. We demonstrated that community sub-divisions of Kenyon cells either aligned with specific sub-types of Kenyon cells (*α/β*-p, *γ*-d, *γ*-m) or compositions of mushroom body neuroblasts (MBp 1-4; Fig. S4).

Together, these analyses underscore the close relationship between the connectivity-based community structure and the functional annotations of neurons. However, they also suggest that this interpretation is inexact; connectivity-based community structure does not always mirror expert-defined neurobiological ontologies, suggesting that there is additional complexity to brain organization that is not entirely captured by current annotation schemes (but also the opposite: that complexity in cell type/function is not fully captured by synaptic connectivity). In sum, these results support the idea that connectivity-driven community analysis and expert annotations provide complementary views of brain function, and together they offer a richer understanding of the organization of the *Drosophila* nervous system.

### A Taxonomy of Community Archetypes

The nested SBM yields a large number of fine-grained communities, each representing a statistically coherent set of neurons with similar connectivity patterns. While these communities capture the microarchitecture of the connectome, their sheer number and heterogeneity make it difficult to identify broader organizational principles. Two communities may be highly similar in their wiring roles yet appear in very different branches of the SBM hierarchy, and conversely, adjacent branches may contain communities that are structurally dissimilar. Thus, the SBM alone does not provide an interpretable mesoscale summary of the types of communities that recur across the brain.

To uncover this higher-order structure, we treated each community as an object with its own anatomical and connectivity signature, characterized by a set of 63 features describing its wiring motifs, spatial footprint, flow properties, and local graph topology. We embedded these feature vectors into a low-dimensional space and clustered them [77] to obtain a set of community archetypes–groups of communities that play similar wiring roles despite differing in size, location, or specific cell types. See Fig. 6 for an overview of this approach, including descriptions of embedding dimensions (principal components). This procedure therefore complements the SBM; the SBM finds the *instances* of communities, while meta-clustering reveals the *classes* of communities that constitute the fundamental organizational motifs of the adult fly brain.

**Figure 6.**
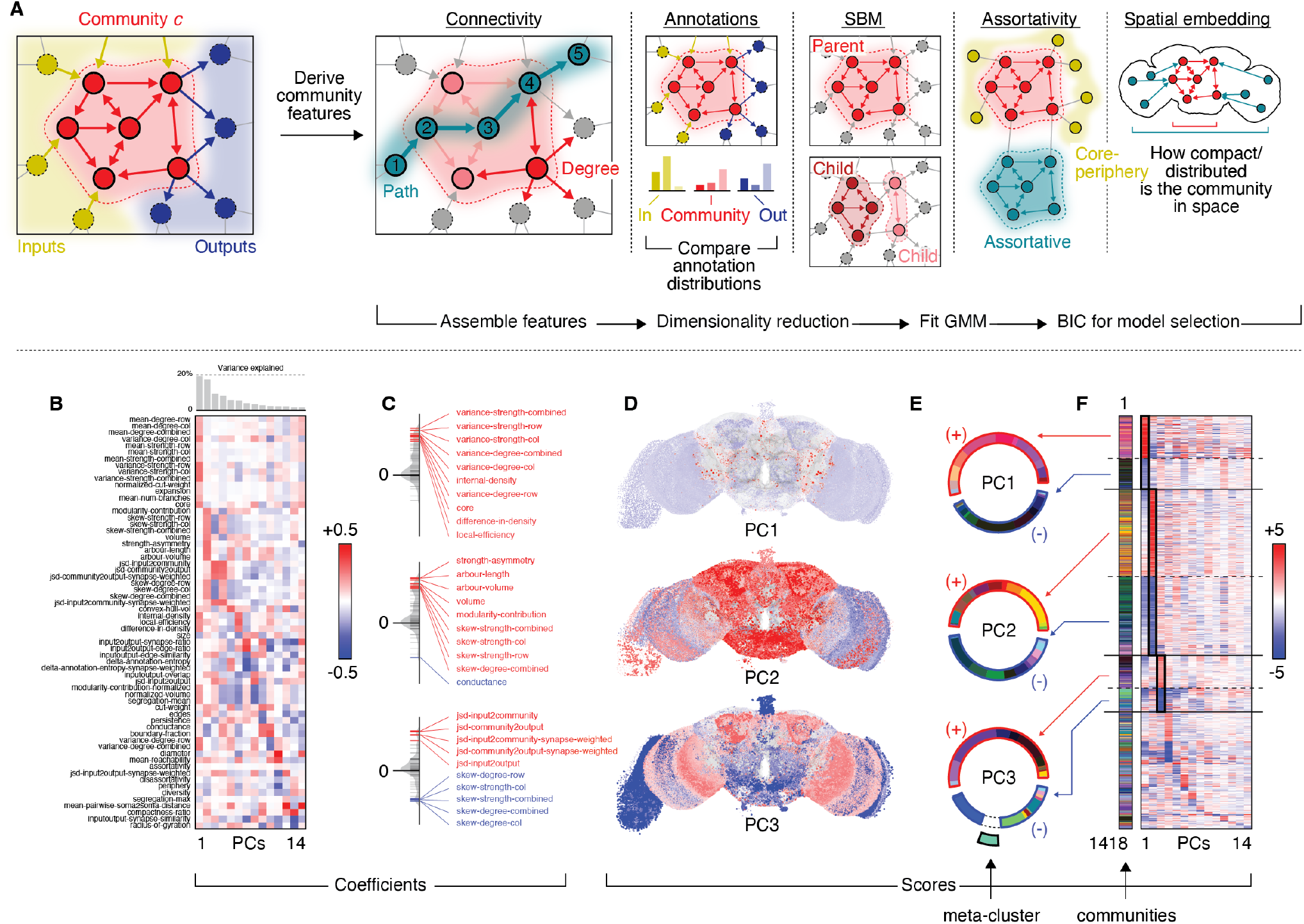
Meta-clustering procedure and principal component analysis of community features. (*A*) We consid-ered communities as separate subgraphs. For each community, we derived a set of features, arranged those features into a community *×* feature matrix (z-scored columns) to which we applied PCA. We clustered communities in feature space using Gaussian mixture models. This figure focuses on those components. PCA returned paired eigenvectors – coefficients and scores – with dimensions feature *×* component and community *×* component, respectively – along with a diagonal matrix of singular values. (*B*) Coefficients for the first 14 components (*>* 85% of variance). (*C*) The distribution of coefficients for the first three components, highlighting the top 10 features by absolute value. (*D*) Component scores mapped to communities and plotted in anatomical space. (*E*) Meta-clusters estimated using the GMM were assigned unique colors. We identified the PC to which each community was maximally affiliated (maximum absolute PC score; see panel *F*). We then calculated the probability distribution of communities with maximal affiliation to the first three PCs, separating positive and negative scores. The aim of this analysis was to give the reader a general sense of which meta-clusters were strongly linked to which PCs and therefore the features shown in panels *B* and *C*.

The resulting taxonomy provides a compact description of mesoscale structure, identifying broad families of communities that differ in their connectivity roles–such as integration modules, feed-forward relays, local recurrent circuits, and wide-field hubs. These archetypes reveal how diverse parts of the brain reuse similar circuit motifs and expose large-scale patterns that are otherwise obscured at the fine granularity of the SBM.

In more detail, Gaussian mixture models grouped fine-scale communities into 45 meta-clusters based on their feature vectors (Fig. S8; Fig. S9). We found that meta-clusters co-localized in both embedding (Fig. 7A) and anatomical space (Fig. 7B,C). The embedding analysis was particularly insightful, as it allowed us to qualitatively compare anatomy with community features. For instance, by aligning the anatomically annotated plot in Fig. 7A with the features shown in Fig. 7D, we learned that communities dominated by visual-centrifugal neurons tended to be internally dense and that those same communities, along with the communities composed of neurons in the central body, also exhibited high levels of local efficiency, suggesting that signals can rapidly propagate from one member of the community to another. A fraction of those same central communities were also materially expensive based on arbour volume, but were nonetheless exceptionally homogeneous in terms of their interactions with other communities, forming mostly segregated, assortative motifs. In contrast, the optic lobe communities participated in more varied interaction motifs. Some of the features aligned poorly with the anatomical labels. For instance, the similarity of incoming and outgoing connectivity profiles did not align neatly with either anatomical annotations or meta-cluster labels. The same is true for the similarity of input-to-output annotations.

**Figure 7.**
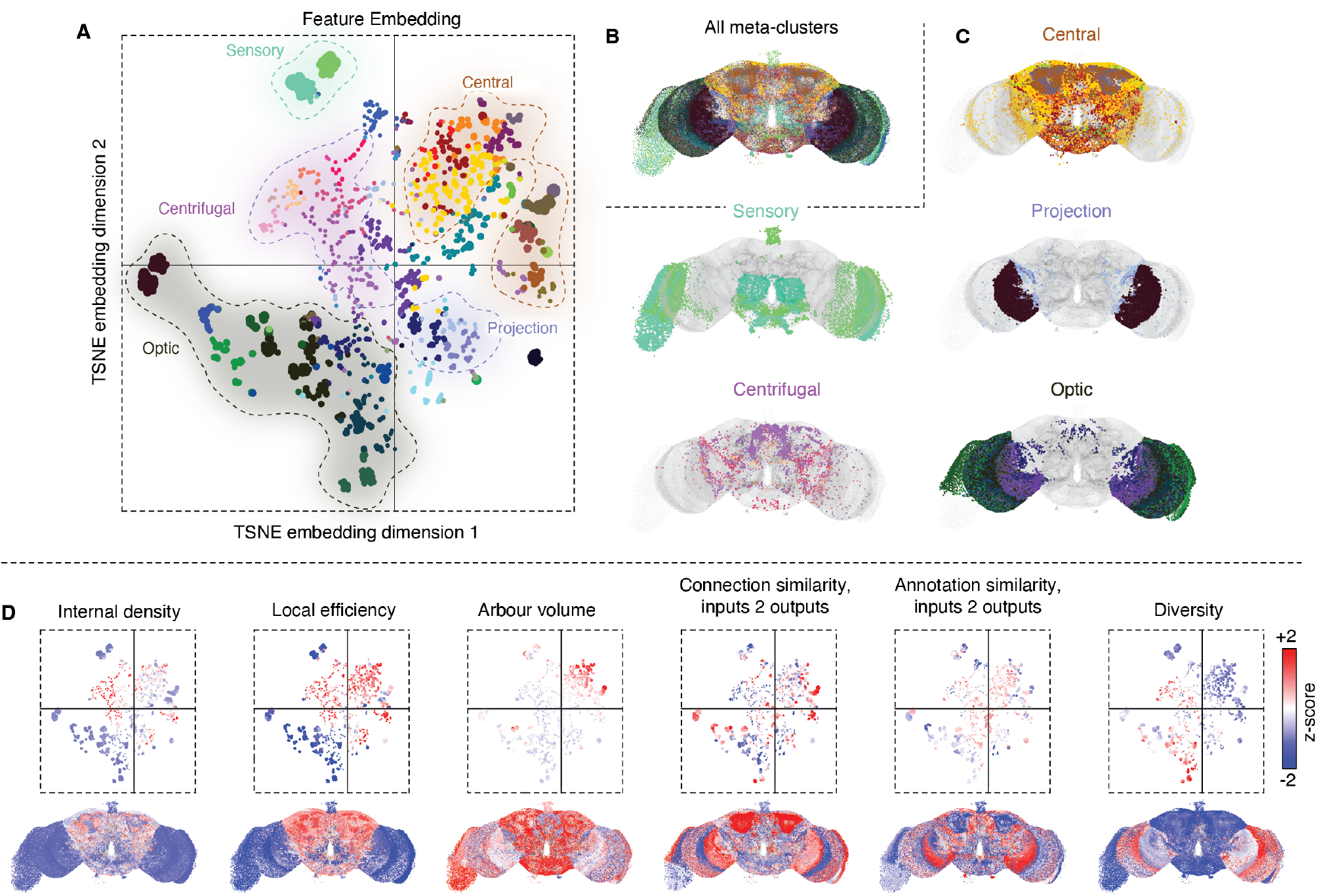
Meta-clusters of communities based on sub-network feature similarity. The consensus meta-clusters were obtained by identifying the partition that was maximally similar, on average, to the partition ensemble corresponding to the top 1% of solutions generated by the GMM based on their BIC. (*A*) To aid in visualization and interpretation, we used *t*-distributed stochastic neighbor embedding (tSNE) to embed the feature matrix in two dimensions (*perplexity* = 20; *exaggeration* = 5). We colored points based on their meta-cluster labels and outlined (broadly) the locations where different `super_class` annotations were concentrated. (*B*) All neurons labeled by their meta-cluster. (*C*) We show meta-clusters that overlapped heavily with several annotations. Note that multiple meta-clusters are visualized in each plot. (*D*) We can also use the tSNE embedding to help visualize the relationships between different meta-clusters, their features, and how those features are localized, anatomically. In this panel, we show a select set of community features in embedding space (top) and in anatomical space (bottom).

To this point, these results suggest that fine-scale communities can be grouped into a relatively small set of structurally distinct archetypes–meta-clusters–based on their community-level features and suggest that the same architectural principles are reused throughout the brain. A question remains: as fine-scale communities are merged into larger communities across the SBM hierarchy, how are features transformed? Do parent communities relate to their children in terms of feature similarity? Do parents preserve the structural signature of their children – i.e. similar feature profiles – or do they aggregate communities into a larger and more dissimilar assembly – i.e. the larger parent is distinct from its children?

To address these questions, we extended our previous analysis by estimating features for communities at *all* levels of the hierarchy. As expected, we found that as hierarchical level increased and communities grew in size, the mean within-community feature similarity tended to decrease, suggesting that larger communities were less homogeneous and increasingly diverse (Fig. 8A). However, the observed levels of similarity were far greater than what would be expected by chance (random groupings of communities into meta-clusters; *p <* 10^−3^), suggesting that homophily–the tendency for similar children to be grouped together–was nonetheless a powerful guiding principle of community organization.

**Figure 8.**
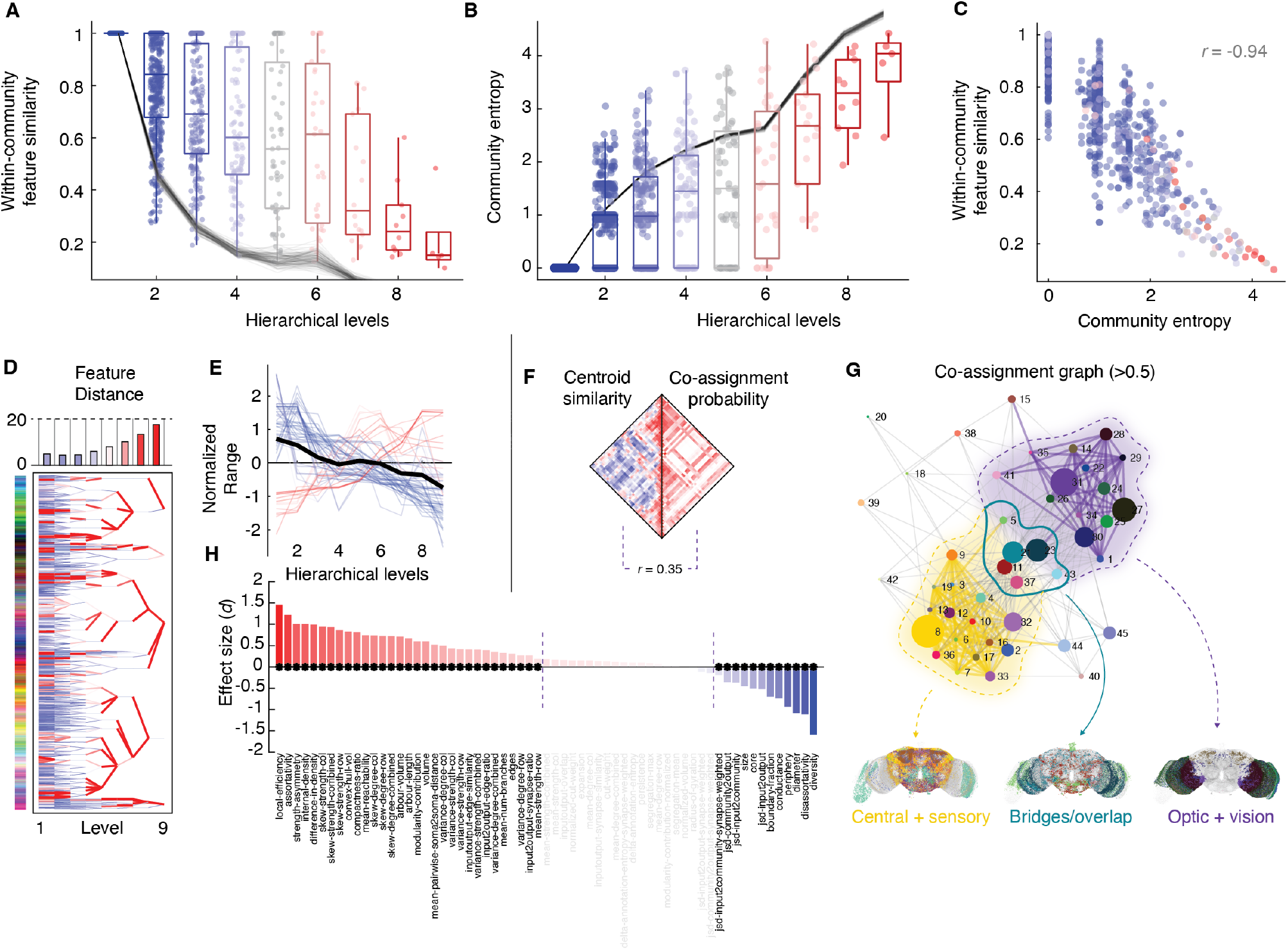
Meta-cluster statistics across hierarchical levels. (*A*) Within-community feature similarity across hierarchical levels. (*B*) Community entropy (based on meta-cluster composition). (*C*) Correlation of community entropy and within-community feature similarity. (*D*) Distance between parent and child feature vectors. Thin blue and thick red lines link similar and dissimilar communities to one another, respectively. (*E*) Normalized ranges for community features as a function of hierarchical level. (*F*) Frequency with which meta-clusters were grouped in the same community (averaged over all levels) (*right*). Meta-cluster centroid similarity (*left*). Correlation between the two matrices (*r* = 0.35). (*G*) The coassignment matrix can be represented as a graph and visualized using spring-embedding layouts to reveal two overlapping groups of meta-clusters: one composed of meta-clusters associated with vision processing and another composed of sensory and central brain neurons. These two groups were interlinked by a third category of bridge meta-clusters. (*G*) We compared the two non-overlapping groups along each of the 63 features. Here, we rank features by effect size. Asterisks indicate statistical significance.

Additionally, we calculated an entropy measure for each community based on its internal composition of meta-clusters. For this analysis we regard fine-scale meta-clusters as atomized and functionally-specific groupings of neurons. Note that here, “function” refers to putative functional roles ascribed to a meta-cluster based on its architectural properties. Intuitively, then, larger communities with high levels of entropy are composed of many different kinds of metaclusters; those with low levels of entropy are internally homogeneous and composed of a relatively few different meta-cluster labels. Indeed, we find entropy increases with community size and hierarchical level (Fig. 8B). However, like within-community feature similarity, the observed entropy levels were much lower than expected by chance (*p <* 10^−3^), reaffirming the central role of homophily in explaining the SBM hierarchy. Indeed, our two primary measurements–within-cluster feature similarity and community entropy–were tightly associated with one another (*r* = −0.94, *p <* 10^−15^; Fig. 8C).

The above analyses described how network-wide measures scale across the hierarchy but tell us little about specific community mergers. That is, do communities with similar features tend to merge with one another? Can we explain why some parent-child community pairs are similar to one another in feature space while others are not? To address this and related questions, we performed the following analysis. First, we used the dendrogram generated by the SBM to assess how similar the feature vectors of each parent and child community were to one another (Fig. 8D). Interestingly, we found that parent-child distance was, on average, small over the first several levels, specifically between children in levels 1, 2, and 3 and their parents in levels 2, 3, and 4, respectively. However, as communities became much larger, parent-child distance grew. More generally, we found that large parent-child feature distances corresponded to cases when there was a large difference in the size of parent and child communities (*r* = 0.45; *ρ* = 0.46) and the overlap in their membership (*r* = − 0.30; *ρ* = − 0.56; Fig. S13). That is, the largest feature transformations occurred when small communities were aggregated into larger communities where they made up only a fraction of the community composition.

Next, we asked how the feature repertoire varied across hierarchical levels. We reasoned that larger ranges corresponded to greater diversity of feature profiles; conversely, smaller ranges corresponded to homogenization of feature similarity. After compiling these statistics for each level, we then z-scored the ranges for each feature, standardizing their units. We found that the range of values decreased monotonically for most features (Fig. 8E). A few measures exhibited increases in their ranges; however, most could be attributed to changes in community size. These outliers included within-community volume and number of binary edges, community size, and arbour length and volume. These observations suggest that the repertoire of functions supported by communities homogenizes as communities merge.

As a final exploratory analysis, we calculated how frequently meta-clusters were coassigned to the same community across all hierarchical levels (Fig. 8F). We found that there was a slight homophilic tendency, in that the more similar meta-clusters were to one another in terms of their features, the more likely they were to be grouped into the same community (*r* = 0.35; *p <* 10^−15^). This effect also reinforces two broad classes of meta-clusters – one containing central and sensory neurons and another associated with optic lobes and vision (Fig. 8G). When we compare these groups in terms of their features, we find that the central+sensory group is highly efficient and assortative, whereas the optic+vision group participates in diverse community motifs and has large diameter (Fig. 8H).

Altogether, these results suggest suggest that fine-scale communities can be grouped into a small set of meta-clusters on the basis of their local features. As communities and meta-clusters get aggregated across the hierarchy, the resulting parent clusters broadly preserve features of their children, but become increasingly homogeneous and less defined.

### Additional analyses

Besides the analyses reported in this section, we also carried out a number of additional analyses whose purposes were not central to the primary aims of this paper. These analyses include: tracking community features along canonical visual pathways (Fig. S6); demonstrating the co-localization of anatomical labels in embedding space (Fig. S7); visualizing the first 10 principal components of community features in embedding space (Fig. S10); and highlighting differences in features across meta-clusters (Fig. S11 and Fig. S12).

## DISCUSSION

Here, we analyze the connectome of an adult female fruit fly. We focus on its community structure. We show that communities are dominantly (but not exclusively) assortative, and that non-assortativity is expressed in communities of neurons involved in visual processing. We show that communities are generally well-aligned with a single annotation (for a given level of annotation). Again, we show that some communities deviate from this principle and are affiliated strongly with multiple annotations. These deviations involve intrinsic neurons in the central brain and sensory neurons associated with mechanosensation and olfaction and multi-sensory integration. Finally, we show that communities are heterogeneous in terms of their local properties and can be grouped into meta-clusters with similar features, suggesting that they may represent functional primitives that can be aggregated into larger, more complex circuits at coarser levels of the hierarchy.

### Most communities interact assortatively

Popular methods for detecting communities feature biases that hinder the detection of nonassortative structure. As a consequence, it is unclear whether the as-sortativity observed in meso-scale connectomes reflects this methodological limitation. Here, we use a stochastic blockmodel to detect communities. The blockmodel can, in principle, detect bipartite and core-periphery structure, if it exists. Despite this, we find overwhelming evidence that the *Drosophila* connectome is organized largely into assortative communities.

On the other hand, we observe a small number of communities that interact nonassortatively. Of note, these communities are composed of laminar neurons in the optic lobes and centrifugal and tangential neurons, which form cores and peripheries (the optic lobe neurons are more peripheral; the centrifugal neurons are more core-like). These collections of neurons are elements of important vision processing pathways and their non-assortativity may reflect the directed flow of visual information into the central brain.

Although the blockmodel is capable of detecting nonassortative communities, those particular motifs appear infrequently, suggesting that the blockmodel, with its relatively high computational cost, could be supplanted in favor of lighter-weight community detection algorithms–e.g. modularity maximization. Indeed, in a recent study of the larval *Drosophila* connectome, we found that modularity maximization recovered similar communities as the blockmodel [23]. However, there are a number of reasons to favor the blockmodeling approach, some technical–e.g. forces users to be explicit about generative processes and is grounded in inferential statistics, making model comparisons straightforward [78]–and others practical–e.g. the nonassortative interactions, though rare, exhibit a high degree of neuroanatomical specificity and, at face value, map onto circuits where feed-forward and core-periphery motifs align with prior knowledge about directed information flows. An important question for future studies is to better understand the advantages and potential trade-offs among community detection algorithms for understanding connectomes; approaches that are technically superior may have little value in neuroscience if they fail to recover community structure that is, neuroscientifically, not meaningful.

### Communities and annotations are closely related

The FlyWire *Drosophila* connectome comes with detailed annotations, imbuing nodes in the connectome graph with additional information and data. Although not wholly independent (the morphology and connectivity may contribute to an evaluator’s decision to ascribe to a neuron a specific annotation), they provide useful meta-data against which we can compare connectivity-driven communities.

Interestingly, we found that communities and annotations were closely related to one another, an observation made previously in the larval fruit fly brain [23]. Although non-causal, this observation suggests that communities and annotations are flip sides of the same coin and are mutually informative of one another.

Nonetheless, we found evidence that annotations and communities diverge. For instance, we observed that communities in the central brain, especially those containing mechanosensory and olfactory neurons, exhibit no clear preference for a single annotation, but overlap with many different annotations, hinting at their poly-functionality.

Additionally, we show that some annotations are poorly represented by communities. That is, those neurons are embedded in communities with differently annotated neurons. However, many of these annotations contained a small number of neurons and may be incompatible with the scale at which communities are detected.

Ultimately, the communities estimated by the SBM – or any community detection algorithm, for that matter – should not, on their own, supersede biology. Community detection algorithms generally only have access to information about (potentially error-prone) connectivity data and optimize objective functions without any neurobiological constraint–i.e. candidate solutions are generally not rejected on the basis of biological plausibility. Further, estimates of community structure are sensitive to choices in algorithm, model structure, and parameters; while there may be technical reasons to prefer one approach over another [78], the final adjudication should be made based on whether an approach yields useful insight into how a system is organized and functions.

In short, community detection is a powerful tool for highlighting statistical regularities in a network’s wiring diagram. Given the relative newness of large, whole-brain, nano-scale connectomes, it is unclear which community detection methods will prove most useful. Anticipating future connectomes that may be orders of magnitude larger than that of *Drosophila*, the most useful algorithms may simply be those that are capable of scaling up to that size while returning noisy (but refinable) community solutions. It is critical, now, that we benchmark and validate algorithmic accuracy using relatively small connectomes, identifying the points where they align with biology. Only then can begin investigating those misalignments and exposing the full potential of community detection for the discovery of novel circuits as part of a dialog that includes theoreticians as well as experimentalists.

### Communities can be aggregated into meta-clusters with similar architectural features

Communities are idealized as groups of functionally related neural elements. In parallel, many studies have speculated that broad categories of brain function might arise from the combination of many small, functionally narrow circuits or communities.

In most studies, the number of communities is orders of magnitude fewer than the number of neural elements. Here, because the original network contains approximately 140,000 elements, the finest hierarchical level partitions neurons into 1449 communities, a number much greater than reported in most previous applications of community detection.

The large number of communities therefore allowed us to study and compare properties of communities to one another (doing so is not as straightforward when the number of communities is small). This also made it possible to further group communities based on their local network, spatial, and annotational properties to discover community archetypes–communities with approximately similar features that are nonetheless distinct from one another.

Our results suggest that, at the finest hierarchy, communities can be grouped into meta-clusters, the largest of which contained hundreds of communities. Interestingly, as these communities are grouped into larger communities, they retain their original properties until approximately the fourth hierarchical level. At that point, however, the properties of fine-scale communities are dissimilar from their parent community and the range of features gets narrower.

Taken together, these observations suggest that there may exist common, reusable organizational blueprints realized at the community level. This assertion is further supported by the strong link between many communities and known annotations, suggesting that the communities align with neurobiological constructs (in many other systems, annotations or meta-data do not share such a strong correspondence with communities [79–81]).

While some of the communities we detected mapped onto known functional pathways – e.g. the L1-ON and L2-OFF pathways – it nonetheless remains unclear to what extent communities fully circumscribe established circuits. Further, there remains other important questions; are communities merely reflecting groupings that are already known – e.g. annotations – or are they (especially those with no clear correspondence to annotations) discovering novel, functionally important groupings? This underscores the need for close collaboration between theoretical and computational research – e.g. responsible for generating the community assignments – with empirical, experimental research and therefore capable to testing functional roles of communities.

Finally, we note that meta-clusters are defined based exclusively on architectural properties of the connectome. While the spatial distributions of the clusters suggest that they exhibit spatial and cell-type specificity, they do not necessarily constitute functional subunits. Ascribing functional import to these clusters requires integration of behavioral and physiological data to probe and verify the functional distinctiveness of meta-clusters and could be a useful direction for future work.

### Novel neuroscientific insights

In this study we examined the community structure of a nanoscale connectome. Community structure, as an architectural feature, has long been studied by network neuroscientists, albeit mostly at the meso/macroscale and using community detection algorithms designed to recover purely assortative communities [20, 30, 82–84] Our study is unique relative to most neuroscience applications in terms of our community detection approach and the unprecedented topological resolution at which the *Drosophila* connectome was mapped. Specifically, we fit nested SBMs to the connectome, which are capable of detecting more general classes of communities– e.g. disassortative, core, periphery–whereas methods like modularity maximization, which are popular in neuroscience, are restricted to assortative communities. Thus, our study joins a growing, but still relatively small, number of studies applying inferential methods to nano-scale connectomes to discover non-assortative communities [26–28, 39–41, 68, 85–87] Indeed, we demonstrate that, although community interactions are overwhelmingly assortative, their deviations from pure assortativity exhibit distinct cellular and spatial profiles.

Additionally, and as noted earlier, most previous studies, especially those applied to meso/macroscale connectomes, have generally resolved brain networks into roughly 10 to 100 communities. Here, we exploit the massive scale of the whole-brain *Drosophila* connectome, returning a hierarchical partition of the brain into 1,449 distinct communities at its finest hierarchical level —representing an increase of two orders of magnitude over the extant literature. This granular lens permits an unmatched view of micro-architectural organization, proving that the structural hierarchy remains highly interpretable even when atomized down to small collections of individual synapses and functional cells.

Rather than simply quantifying community boundaries, we used this enhanced resolution to systematically quantify properties of the communities themselves. Rather than treating communities as monolithic/homogenous entities, we characterized their high-dimensional feature spaces, integrating multi-modal properties, including spatial embedding, structural connection profiles, and cellular annotations. We used the features as input to a novel, data-driven clustering framework to reveal a hidden layer of organization: a taxonomy of meta-clusters, which we speculate represent latent archetypes or structural “primitives” that could serve as modular building blocks from which the whole brain is wired. By identifying these archetypes, we show that while individual fine-scale communities are highly heterogeneous in their anatomical positioning and specific partners, they converge onto a remarkably low-dimensional set of topological rules. Crucially, this meta-clustering uncovers a clear dichotomy between highly assortative, efficient circuits characteristic of central and sensory pathways and non-assortative, high-diameter processing motifs concentrated within the visual system and optic lobes. This approach provides a repeatable blueprint for digesting nanoscale, synapse-resolution graphs into biologically coherent principles of design [88–90].

### Bridging scales in neuroscience

To better emphasize the relevance of our findings relative to mammalian connectomics and large-scale brain network studies, we note that the significance of the present work lies less in establishing a direct correspondence between Drosophila and mammalian connectivity patterns, and more in the identification of organizational principles that may be universal and shared across both species and spatial scales. While the architecture of Drosophila communities is shaped by species-specific constraints, including brain size, developmental bias, and behavioral/cognitive repertoire, several of the organizational principles examined here, such as hierarchical organization [91] and coexistence of assortative [30] and non-assortative communities [39, 41], have also been observed in mammalian connectomes and largescale brain networks. From this perspective, principles that appear in systems separated by hundreds of millions of years of evolution and orders of magnitude in size become candidates for general principles of neural organization, rather than species-specific properties. Community structure may represent one such principle, with the observed correspondence between network communities and functional specialization reflecting an aspect of nervous systems organization conserved across species.

### Limitations and future work

There are a number of limitations associated with this study that should be considered when interpreting the results. FAt the meso-scale, community structure irst, this is, in essence, an *N* of 1 study; we examined a *Drosophila* connectome from a single individual. While the level of stereotypy across flies is likely greater than that of the mammalian brain, our study does not account for individual variation. Moreover, our study represents a snapshot of that individual at a single developmental time point, making it impossible to assess whether some of the communities described here, if they are indeed close to “ground truth” for this instant, will dissolve or persist over time. Even this snapshot is incomplete; this connectome reconstruction does not include any gap junction, omitting this important mode of neuron-to-neuron communication from our analyses completely.

Second, in time since this study began, two new processed *Drosophila* datasets became available [44, 92], both of which include neurons in the nerve cord (analogous to the spinal cord in vertebrates). Omitting these neurons from our analysis runs the risk of ignoring functionally important interactions with effectors and body segments.

Third, our meta-cluster analysis has some practical limitations. Namely, it forces us to specify a set of features used for the clustering procedure; in principle, different sets of clusters could yield different clustering solutions. To partly mitigate this issue, our clustering is based on principal components, which addresses issues related to correlated features, but does not address the potential issue that the inclusion of an additional feature could yield a different cluster solutions. In general, solving this is issue is not simple – the large space of possible features virtually guarantees that some sub-spaces will not be sampled densely. To this end, many of the features we measured were generally well-established in network science literature, and therefore may be a reasonable sample of measurements. Nonetheless, future work may find it useful to investigate alternative measures, especially those that are uncorrelated with those reported here [7].

The meta-cluster analysis (essentially clustering the output of one clustering algorithm with another) presents some conceptual challenges. Could we have obtained comparable results without the two-step algorithm–e.g. by defining features for individual neurons and clustering them using mixture modeling? While this, in principle, is a perfectly reasonable approach, it misses the point of the meta-clustering analysis. Specifically, we hypothesized that network functions are carried out not by individual neurons, but circuits and sub-networks. In fact, many of the features we used to group communities into meta-clusters are not well-defined for individual neurons; they are properties only associated with groups of neurons. For this reason, it is unlikely that feature-based clustering of individual neurons will yield partitions comparable to our meta-clusters.

Relatedly, the results of the meta-cluster analysis depend on the set of features used to estimate the principal components. Would we obtain similar meta-clusters if we omitted some of the features or included new features? We speculate that these clusters are relatively stable against small perturbations. Our reasoning is two-fold. On one hand, prior work has shown that many graph-theoretic measures are highly correlated, with one another [93]. Therefore, we anticipate that new features (particularly those derived from connectivity) are likely to be correlated with one or more of the existing features, leaving the PCs relatively unchanged. On the other hand, we can estimate the stability of our PCs (the inputs to the clustering algorithm) after excluding sub-sets of features; if the PCs largely unchanged, then we expect the clustering algorithm to return comparable solutions. We found that even after removing a significant fraction of features (30 of the 63), the original PCs were mostly preserved (Fig. S14). These results suggest that the meta-cluster labels are likely to be stable under modest perturbations. However, it is unclear what impact introducing/removing large numbers of features might have on the meta-clustering results. Relatedly, it is unclear what would happen if a small number of features (selected to be near-orthogonal to those used here) were introduced. We leave this exercise for future studies.

As an additional limitation, we point out some of the issues related to our use of the stochastic block-model. Although obtaining high-quality estimates of nested communities, even for a large network containing 140k nodes, can be easily generated locally on personal computers. However, for the purposes of sampling communities from the posterior distribution – the preferred approach over accepting a point estimate – takes much longer (on the order of days on a personal computer, but likely much faster on a high performance computing cluster). Accordingly, and despite their utility for small networks, nested stochastic blockmodels might not be an optimal approach for future, anticipated datasets with millions or billions of neurons. Future studies may benefit from exploring alternative approaches. One possible alternative that has excellent scalability are random dot product graphs (RDPG), embed networks into low-dimensional spaces and rely on scalable machine learning techniques–e.g. Gaussian mixture models–for obtaining clusters. Part of the scalability of this approach is derived from the fact that connectomes at the nanoscale appear sparse, and therefore amenable to dimension reduction techniques (singular value decomposition) for obtaining embedding coordinates. Importantly, RDPGs can be viewed as generalizations of the SBM (the SBM is equivalent to a RDPG where each node in the same community occupies the same position in embedding space) but can also recover continuous relationships between communities – i.e. overlap or partial affiliation with multiple communities – potentially making RDPGs essential tools for nanoscale connectomics [42, 94].

Fifth, an important potential limitation is related to space. In our analyses, we found that most communities were more spatially compact than we would expect given their size and number of neurons. Elsewhere, it has been established that spatial embedding is a key constraint on connectome organization, with distant brain areas forming connections less frequently and, when the do, with weaker connection weights [11, 56, 57, 95]. A version of this rule must also apply to nanoscale connectomes – synapses require spatial proximity, so neurons that never come into close contact with one another can never synapse onto one another [96]. Irrespective of the precise form, this presents a challenge for model-based inference of connectome communities; models that fail to account for spatial proximity may falsely group densely packed clusters of neurons, interpreting their reciprocal connections as evidence that they are a community, but in reality, those connections reflect geometric wiring rules. On the other hand, accounting for spatial relationships (treating it as a nuisance variable to remove from this dataset) may be misguided; evolution tends to co-opt constraints. Through this lens, we should expect that functionally related groups of neurons are co-localized in space, preserving functionality at a relatively low material/metabolic cost [97]. Nonetheless, future work should extend approaches like SBMs (or RDPGs) so that they explicitly include spatial information in the generative process.

Another potential limitation and opportunity for extending our work concerns the meta-cluster analysis. As part of this analysis, we group communities (and by extension, their constituent neurons) into meta-clusters based on their feature similarity. Could we have obtained similar divisions of neurons into groups without the “intermediate” step of first estimating communities? Allowing for some speculation, we consider this possibility unlikely. The primary rationale is that many of the features used to obtain meta-clusters are not defined for individual neurons, e.g. local efficiency and the community interaction metrics (assortativity, coreness, periphery, disassortativity, diversity) are only defined for sub-graphs.

Finally, there has been considerable effort to link biological neural networks with artificial networks – for instance, see [98–101] These types of models, which often necessitate solving or simulating data from a massive system of coupled differential equations, might benefit greatly from an initial dimension reduction step. Rather than arbitrarily coarse-graining the connectome, treating communities as the units of interest would serve to speed up compute times while approximately preserving the connectivity patterns of the underlying neurons.

## MATERIALS AND METHODS

### Drosophila connectome dataset

All connectome data, including synaptic connectivity, three-dimensional coordinates, neuropil volumes, and annotations were obtained from https://codex.flywire.ai/. These derivatives represent the output of a process beginning with nanometer-resolution electron microscopy images of an adult female *Drosophila* [102]. The FlyWire project allowed citizen scientists to trace neurons [103] to reconstruct the connectome. Combined with automated approaches [104, 105], the end result was a complete map of synaptic connections among approximately 139,000 neurons.

Neurons in *Drosophila* are polyadic, thus the same pair of neurons can synapse onto one another at multiple distinct sites. The https://codex.flywire.ai/ are shared in two versions: one in which pairs of neurons were connected if they were linked by, at minimum, five synapses and another in which even a single synapse was considered evidence that two neurons were connected (unthresholded). We opted to analyze the unthresholded connectome, which preserved all synapses.

Annotation data were obtained from at the same time as the connectome data. Annotations were grouped into four distinct categories [106]. The first two annotations are “dense,” in the sense that every neuron is assigned a label. The “Flow” included afferent, efferent, and intrinsic labels. “Super-class” included sensory (periphery to brain), motor (brain to periphery), endocrine (brain to corpora allata/cardiaca), ascending (ventral nerve cord (VNC) to brain), descending (brain to VNC), visual projection (optic lobes to central brain), visual centrifugal (central brain to optic lobes), or intrinsic to the optic lobes or the central brain.

The other three categories are not dense–i.e. not all neurons are assigned a label. The “Class” and “Sub-class” annotations are nested hierarchically within the the parent “Super-class” and describe cell groups reported in the extant literature, e.g. neurons in the optic lobe [107]. The “Cell-type” annotations were, similarly, not dense.

### Community detection

Many studies have shown that biological neural networks at varying spatial scales can be meaningfully partitioned into clusters or communities [10, 21, 23, 24, 27, 36, 108–111] In general, community structure is unknown ahead of time and must be estimated or inferred algorithmically. The procedure is referred to as community detection [14], and there exist many approaches for doing so [38, 94, 112–115] each of which makes specific assumptions about what constitutes a community.

One of the oldest approaches for community detection is the stochastic blockmodel [116–120]. Briefly, blockmodels imagine that every node in the network is assigned a label. The probability that any pair of nodes are connected and the weight of that connection depends only upon the communities to which they are assigned. In general, blockmodels estimate community labels and connection probability/weight distributions so as to maximize the likelihood that the model generated the observed network.

Recently, there has been renewed interest in the application of blockmodels to brain network data [27, 39, 41, 47, 68, 79, 121–124] Here, we leverage a nested and weighted variant of the classical blockmodel to partition the *Drosophila* connectome into hierarchical communities [48]. In our model, we included a term that corrected for nodes’ degrees and assumed that edge weights were drawn from a geometric Poisson distribution (initially we tested alternative distributions and found that they resulted in solutions corresponding to greater minimal description lengths; consequently we moved forward with the geometric Poisson distribution). To fit the parameters of this model, we used a Monte Carlo Markov Chains approach. Briefly, we used a merge/split algorithm to equilibrate the Markov Chain [124]. Following equilibration, we sampled 10000 high-quality partitions, skipping every 10^th^ iteration to mitigate strong serial correlation in our samples. From this ensemble of partitions, we obtained a consensus hierarchical partition [125] that corresponded to the modal community assignment of each neuron. All analyses using communities were carried out on the resulting partition.

### Annotation enrichment analysis

Throughout this manuscript we describe an “enrichment” procedure for linking continuous data to categorical labels–e.g. annotations. The aim of this procedure, in more detail, was to determine if a particular variable was over-expressed (enriched) within a given annotation category. To do this, we calculated the variable of interest’s mean value within all neurons associated with a specific annotation. We then randomly permuted the annotation labels (keeping the total number constant) and calculated the new mean. We repeated this procedure 1000 times, generated a null distribution of mean values. The enrichment score for that variable within that annotation category is the original mean expressed as a z-score with respect to the null distribution.

### Community interaction motifs

The SBM is sensitive to both assortative and nonassortative community configurations. To identify these types of interactions, we considered community dyads – i.e. pairs of communities, *r* and *s*. We classified their interaction based on the density of connections – recurrent densities *d*_*rr*_ and *d*_*ss*_ as well the block of directed connections from *r* to *s, d*_*rs*_. For each dyad, we labeled its interaction as assortative if both *d*_*rr*_ *> d*_*rs*_ and *d*_*ss*_ *> d*_*rs*_; it was labeled as disassortative if *d*_*rs*_ *> d*_*rr*_ and *d*_*rs*_ *> d*_*ss*_; *r* was a core and *s* a periphery if *d*_*rr*_ *> d*_*rs*_ *> d*_*ss*_ and *vice versa*.

The dyadic labels were stored in community *×* community matrices. For each community, we obtained an motif type count by calculating the number of assortative, disassortative, core, peripheral, and nonassortative (disassortative + core + periphery) dyads that it participated in. We also calculated a diversity score for each community as the entropy of the probability distribution, [*p*_*assortative*_, *p*_*disassortative*_, *p*_*core*_, *p*_*periphery*_].

Note that some interactions were considered “ambiguous”. The interaction motif would be classified as ambiguous if *d*_*rr*_ = 2 but *d*_*ss*_ = *d*_*rs*_ = 1. Had *d*_*ss*_ = 2 and *d*_*rs*_ = 1, then the interaction would have been labeled assortative. Had *d*_*ss*_ = 0 and *d*_*rs*_ = 1, then the interaction would have been labeled as core-periphery. However, the tie (*d*_*ss*_ = *d*_*rs*_) makes it impossible to unambiguously classify this interaction.

This type of ambiguity occurs only when synapse densities (the # of synapses divided by the number of neurons in two groups) are exactly equal. Fortunately, equal densities are extremely rare and are most common when motifs involve very small communities composed of a handful of neurons. Because the global assortativity/core/periphery/disassortative scores are weighted by community size, these rare ambiguous motifs are further downweighted and contribute minimally to our results.

### Community meta-cluster analysis

The SBM generated a set of nested community labels. Each community sub-graph can be analyzed separately. Our focus was on the fine-scale communities, but we also calculated this set of measures for all communities, irrespective of their size or level in the hierarchy. Here, we describe the set of measures.

- **Modularity contribution (1)**: The contribution to the modularity quality function, *Q*, by community : 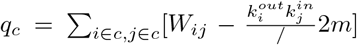, where 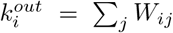 and 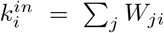 and 2*m* = Σ_*i*_ *k*_*i*_.
- **Normalized modularity contribution (2)**: Modularity contribution, *q*_*c*_, is biased towards larger communities. Here, we normalize *q*_*c*_, dividing it by *n*_*c*_(*n*_*c*_ − 1), where *n*_*c*_ is the number of neurons assigned to community *c* [38].
- **In/out/combined degree (moments) (3-11)**: For each neuron, we calculated its in/out degree as well as its combined in + out degree. We identified the neurons assigned to each communities and, given the distribution of their degrees, calculated its mean, variance, and skewness. This resulted in 9 separate measures (in/out/combined *×* mean/variance/skewness).
- **In/out/combined strength (moments) (12-20)**: Identical set of calculations, but for in/out/combined strength (weighted degree).
- **Community size (21)**: Number of neurons assigned to a community, *n*_*c*_.
- **Edge count (22)**: The number of edges (binary) that fall within a community.
- **Volume (23)**: The number of synapses (weighted) that fall within a community.
- **Strength asymmetry (24)**: For each neuron in community *c*, we calculate the sum of squared differences in neurons incoming and out-going weighted degree 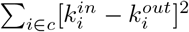.
- **Density (25)**: The number of edges (binary) that fall within a community normalized by total number of possible connections.
- **Normalized volume (26)**: The number of synapses (weighted) that fall within a community normalized by total number of possible connections.
- **Local efficiency (27)**: Given only a community subgraph, we calculate the path length between all pairs of neurons that comprise that community (excluding the rest of the network). We calculate local efficiency as: 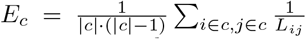 That is, local efficiency measures the mean reciprocal shortest path length between neurons within that community [126].
- **Diameter (28)**: The length of the longest shortest path between pairs of nodes assigned to community *c*.
- **Mean reachability (29)**: It is possible that a community subgraph includes disconnected neurons, such that there exists neurons in the community that are not reachable from other members of the community–i.e. *L*_*ij*_ = ∞. Mean reachability is the fraction of non-infinite path lengths among all pairs of neurons belonging to the community.
- **Cut weight (30)**: Total number of synapses (weighted connections) between community *c* and all other communities [127].
- **Normalized cut weight (31)**: Cut weight can be biased by size of communities. This measure normalizes by the size of the two communities.
- **Conductance (32)**: Similar to cut weight, but normalized by nodes’ strengths [128, 129].
- **Expansion (33)**: Similar to cut weight, but measures the mean number of synapses leaving community *c* per node.
- **Boundary fraction (34)**: The proportion of connections from community *c* that leave the community.
- **Input-to-output synapse ratio (35)**: We identify all incoming and outgoing connections to community *c*. We count the total number of incoming and outgoing synapses. We express these numbers as a ratio.
- **Input-to-output edge ratio (36**: Identical to “Input-to-output synapses ratio” but ignores edge weights; ratio of the number of incoming binary connections to outgoing binary connections.
- **Input-to-output synapse similarity (37)**: We identify all pre- and post-synaptic neurons. From these, we calculate the mean connectivity profile of pre-/post-synaptic neurons. This measure is the cosine similarity of those vectors.
- **Input-to-output edge similarity (38)**: Similar to the previous measure, we identify all pre- and post-synaptic neurons. From these, we calculate the mean binary connectivity profile of pre-/post-synaptic neurons. This measure is the cosine similarity of those vectors.
- **Input-to-output overlap (39)**: We identify the sets of pre-/post-synaptic neurons and calculate the Jaccard similarity of these two sets.
- **Segregation (maximum) (40):** Within-community connection density for community *c* minus the maximum between-community connection density across all other communities.
- **Segregation (mean) (41):** Within-community connection density for community *c* minus the mean between-community connection density across all other communities.
- **Jensen-Shannon Divergence (input-to-output annotations) (42):** Let *P* and *Q* be the distributions of annotations of all neurons that send inputs or receive outputs from community *c*, respectively. We can measure how similar those annotations are to one another with the Jensen-Shannon divergence: 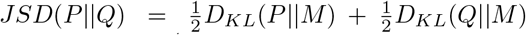, where 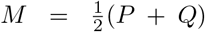 and 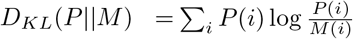. The JSD does not satisfy the triangle inequality and is therefore not a proper distance metric. However, 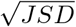 does. We use this formulation here. Intuitively, this measure tells us how similar populations of pre-/post-synaptic neurons are to one another in terms of their annotations.
- **Jensen-Shannon Divergence (input-to-community annotations) (43):** Identical to the previous measure, but where *P* and *Q* correspond to pre-synaptic neurons and the neurons that make up community *c* [130].
- **Jensen-Shannon Divergence (communityto-output annotations) (44):** Identical to the previous measure, but where *P* and *Q* correspond to post-synaptic neurons and the neurons that make up community *c*.
- **ΔAnnotation entropy (45):** We wanted to know whether there was a big increase or decrease in the diversity of input-to-output annotations. That is, does a community behave like a bottleneck, receiving inputs from diversely-annotations neurons before delivering them to a set of narrowly-annotated neurons? Or does it behave like a fan, receiving inputs from narrowly-annotated neurons and delivering them to a diversely-annotated neurons. If *P* and *Q* annotation distributions for pre-/post-synaptic *P* neurons, respectively, then we calculate *H*_*P*_ = Σ_*i*_ *P*(*i*) log *P*_*i*_ and *H*_*Q*_ = Σ _*i*_ *Q*(*i*) log *Q*_*i*_. We then calculate the difference Δ*H* = *H*_*P*_ ™ *H*_*Q*_. When Δ*H* > 0, the community behaves like a bottleneck; when Δ*H* < 0 it behaves more like a fan.
- **Jensen-Shannon Divergence (input-to-output annotations, synapse weighted) (46):** The above JSD measures identified pre-/post-synaptic sets of neurons. In doing so, they ignored synaptic weights. Here, we calculate annotation distributions, *P* and *Q*, but rescale contributions by synaptic weight. To illustrate the difference, consider neuron *p* that makes 10 pre-synaptic synapses onto members of community *c*. In the previous measures, the annotation of this neuron would be counted once when calculating *P*. Here, it gets counted 10 times (weighted by the number of synapses).
- **Jensen-Shannon Divergence (input-to-community annotations, synapse weighted) (47)**: Same as above, but for input and community neurons.
- **Jensen-Shannon Divergence (community-to-output annotations, synapse weighted) (48)**: Same as above, but for community and output neurons.
- Δ**Annotation entropy, synapse weighted (49)**: Same as the unweighted ΔAnnotation entropy, but weighted by synapse count.
- **Community persistence (50)**: Jaccard similarity between parent and child communities.
- Δ**density (51)**: Difference in density between parent and child communities.
- **Assortativity (52)**: Fraction of interactions classified as assortative.
- **Disassortativity (53)**: Fraction of interactions classified as disassortative.
- **Core (54)**: Fraction of interactions classified as core.
- **Periphery (55)**: Fraction of interactions classified as periphery.
- **Diversity (56)**: The entropy of the distribution {Pr(*Assortative*), Pr(*Disssortative*), Pr(*Core*), Pr(*P*treating)}
- **Radius of gyration (57)**: Mean squared Euclidean distance of all neurons in community *c* to their center of mass.
- **Mean pairwise soma-to-soma distance (58)**: Mean pairwise Euclidean distance between all neurons assigned to community *c*.
- **Convex hull volume (59)**: Volume of the convex hull around all neurons in community *c* and their complete arbours.
- **Compactness ratio (60)**: Ratio between the convex hull volume and a sphere with the same radius of gyration as community *c*.
- **Mean number of branches (61)**: For all neurons assigned to community *c*, the average number of branch points in their arbours.
- **Arbour length (62)**: The total length of arbourization (ignoring cylinder width).
- **Arbour volume (63)**: Same as above, but where each line segment was treated as a cylinder whose radius was equal to the the arbour thickness.

Some of the above measures are not defined for singleton communities. Due to this limitation, we omitted 31 singleton communities from analysis. For each measure, we then standardized (z-scored) its value across communities all 1418 remaining communities. The result was a 1418 *×* 63 feature matrix. This matrix was used as input to fit a Gaussian mixture model. Briefly, Gaussian mixture models are generative models of multivariate data. They posit that data are drawn from multivariate Gaussians; the fitting procedure involves determining the number and parameters of these Gaussians. We varied the number of Gaussians from *k* = 2 to *k* = 250, running the optimization algorithm 100 times at each *k*. We also calculated the Bayesian information criterion (BIC) for each solution. This measure assigns models a scores based on how well they fit the observed data while penalizing for model complexity (as complex models with many parameters will tend to outperform models with few parameters).

We found that the optimal number of components was between 35 and 43 with a mode of 39 (Fig. S8a). We retained the partitions associated with the top 1%, *eriphry* them as a representative sample of “good” clusters (Fig. S8b). Rather than describe features of the ensemble in a statistical sense, we estimated consensus clusters–i.e. the partition that was maximally similar, on average, to the partitions that comprised the ensemble. As a measure of similarity, we used the Adjusted Rand Index (ARI; Fig. S8c). We used a zero-temperature simulated annealing algorithm to optimize mean ARI, yielding an optimal partition of 45 metaclusters. As expected, the meta-clusters were largely aligned with variation in PCs (Fig. 6e,f; see Fig. S9 for depictions of all 45

## Author contributions

- Conceptualization: RB, MGP, CS
- Data curation: RB
- Formal analysis: RB
- Investigation: RB, OS, MGP, CS
- Methodology: RB, OS, MGP, CS
- Software: RB
- Visualization: RB
- Writing - original draft: RB, ODR, NL, MGP, CS
- Writing - review & editing: RB, ODR, NL, SD, BL, CWL, BM, LP, OS, BTC

## Data and code availability statement

Original data are publicly available through https://codex.flywire.ai/ with no restrictions. The Fly-Wire derivatives used as part of this manuscript are available here: https://zenodo.org/records/20399924. Code for performing community detection, running all primary analyses, and generating Figures 2-8 is available here: https://zenodo.org/records/20649529.

---

**Figure S1.**
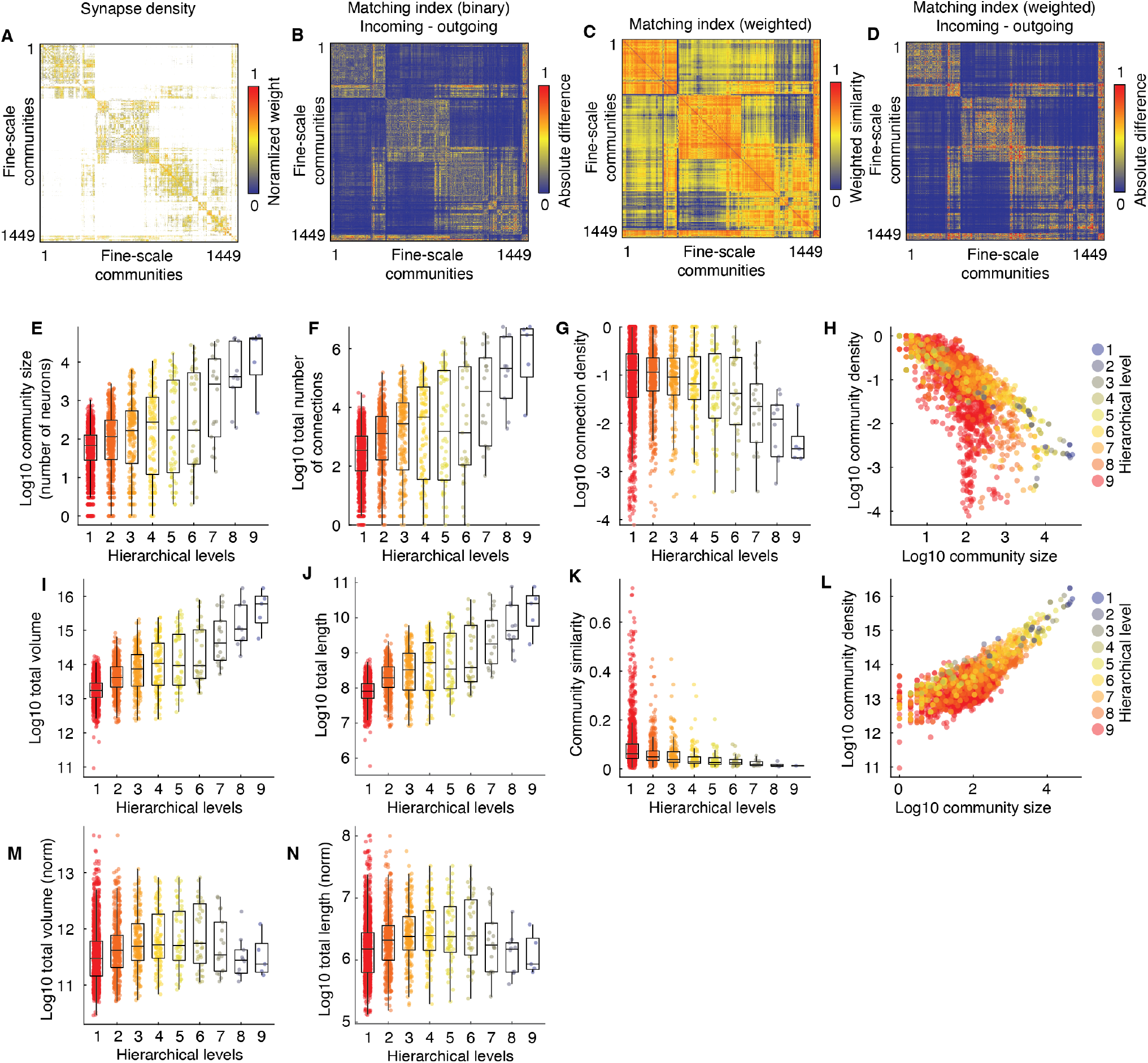
Additional community statistics. (*a*) Connectivity matrix coarse-grained by fine-scale communities. That is, each row/column represents one of 1449 communities. Weights correspond to synaptic density. For communities *r* and *s*, synaptic density is defined as 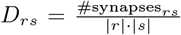, where |*r*| is the size of community *r*.(*b*) Difference in matching indices estimated using only incoming *versus* outgoing connectivity profiles. (*c*) Matching index estimated using weighted information. (*d*) Difference in between incoming/outgoing weighted matching index. Panels *e*-*n* show community statistics as a function of hierarchical level.

**Figure S2.**
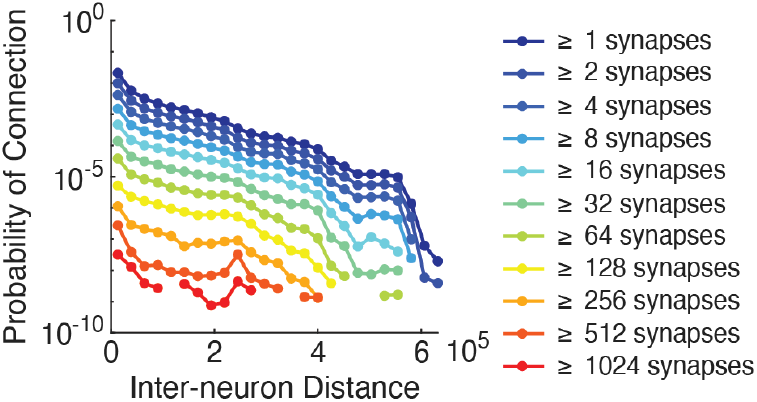
Effect of inter-soma distance on connection probability. For pairs of neurons, we calculated their intersoma distance. We then binned neuron pairs by distance and, within each bin, calculated the probability that neurons in that bin were connected by *K* synapses. Here, we plot the log probability as a function of distance.

**Figure S3.**
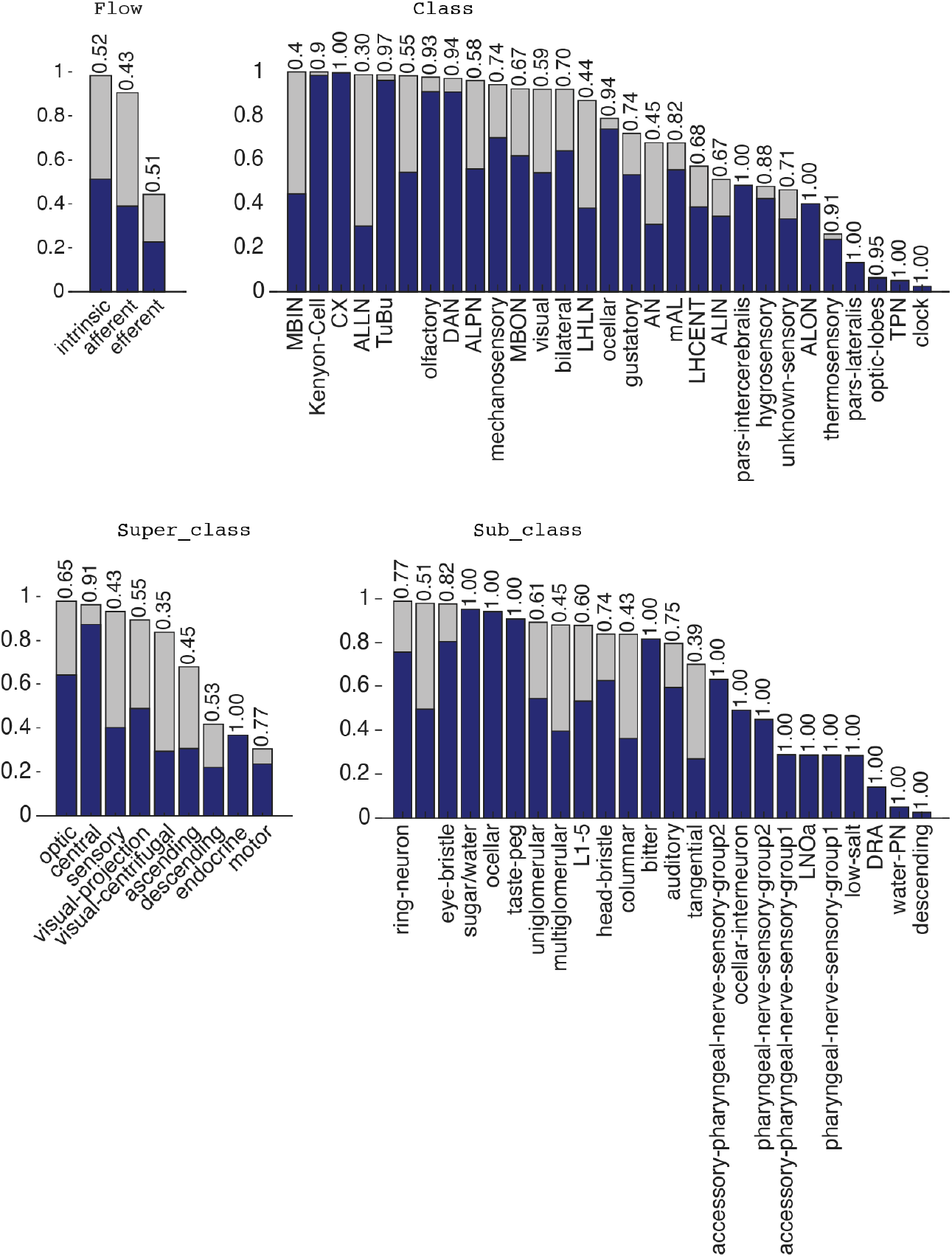
Comparing optimized, multi-community Dice coefficients with multi-scale, single-community coefficients. In the main text, we used an optimization algorithm to combine fine-scale communities so as to maximize the overlap with respect to neuronal annotations. Here, we compare each annotation’s optimized overlap score (Dice coefficient) against its best single-community overlap score obtained at any level of the hierarchy.

**Figure S4.**
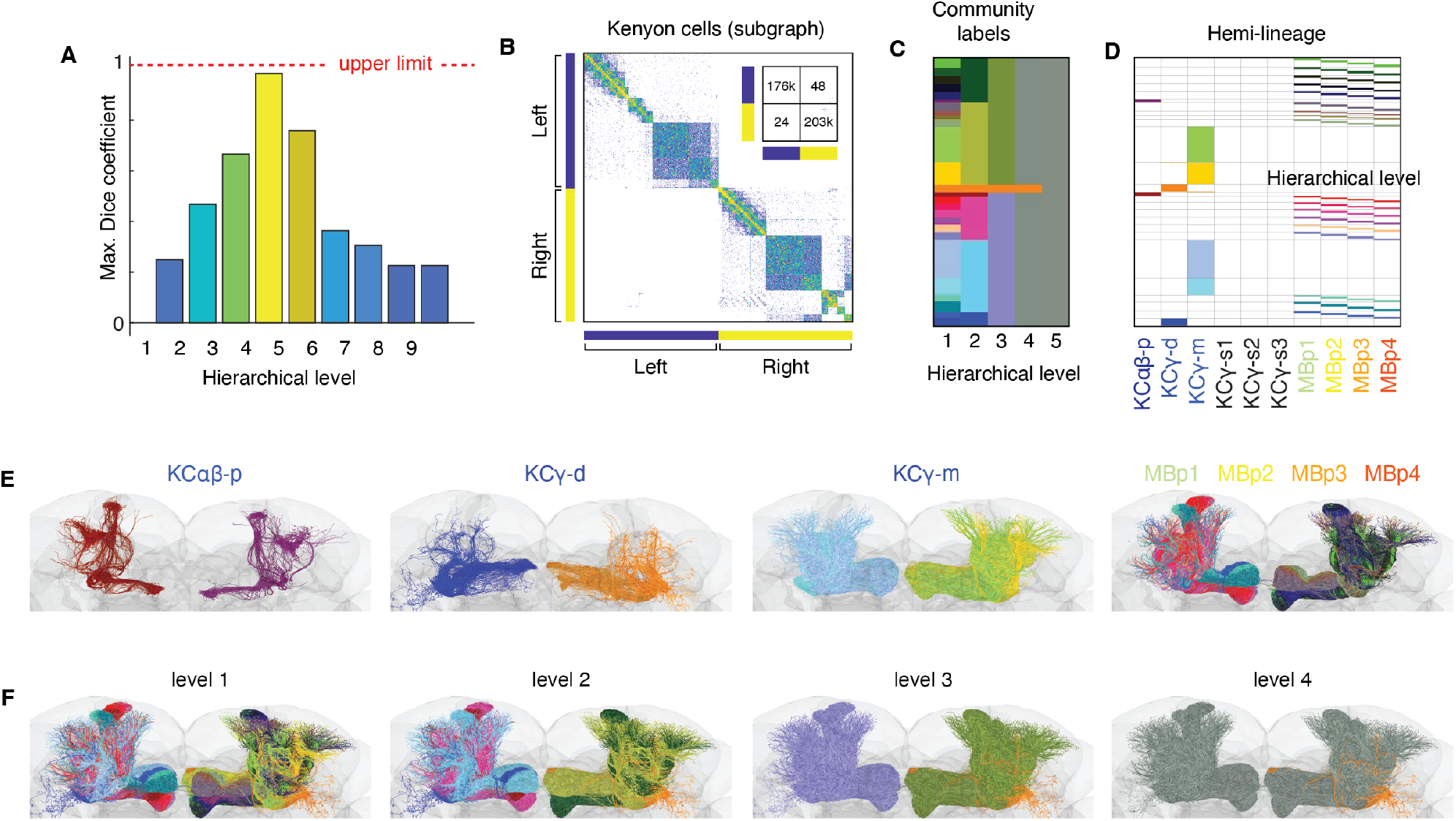
Hemilineage analysis of Kenyon cell communities. (*a*) Maximum Dice coefficient of Kenyon Cells (KCs) annotatio nwith communities at different hierarchical levels. (*b*) Subgraph of KCs ordered by community with side (left or right) labeled. The inset shows the number of synapses between neurons on either side of the brain. (*c*) Community labels shown across the first five hierarchical levels. Note that to visually emphasize different communities, we reassigned communities new colors for this figure; this colormap is not the same as the one used in the main text for community labeling. (*d*) Hemilineage labels for each neuron colored by the community to which that neuron was assigned. (*e*) Anatomical view of the different KC types. (*f*) Anatomical plot showing KCs at different levels of the hierarchy.

**Figure S5.**
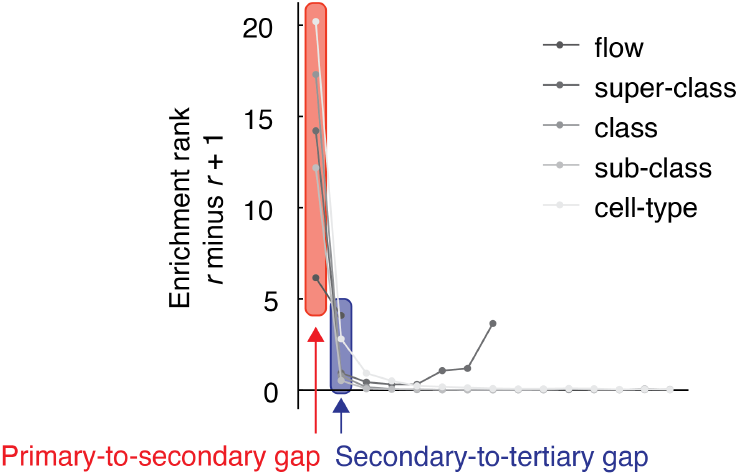
Enrichment gap. For each community, we identified the annotation for which it was maximally enriched and retained that number. We compared that “primary” enrichment score to the score of the annotation with the second greatest enrichment score. The difference between those scores is the “primary-to-secondary” enrichment gap. We repeated this measurement for secondary and tertiary scores, tertiary to quaternary, quaternary and quinary, and so on. We found that the primary-to-secondary gap was, on average, the greatest.

**Figure S6.**
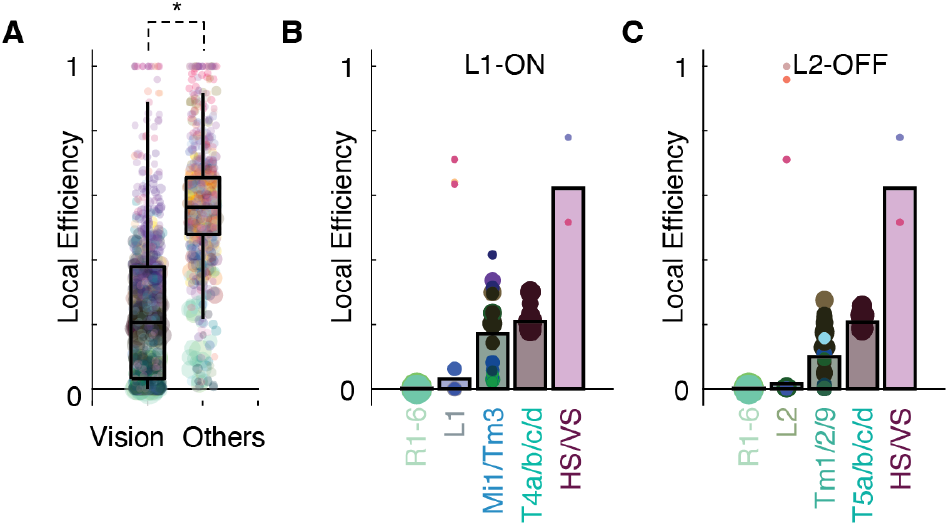
Community features vary along canonical ON/OFF visual pathways. We tracked local efficiency along two established visual pathways – the L1-ON and L2-OFF pathways. We found that, progressing from photoreceptors to vertical/horizontal cells, local efficiency of communities monotonically increased. This increase mirrors a more general trend in which communities composed of neurons strongly associated with vision processing exhibited reduced efficiency relative to all other neurons.

**Figure S7.**
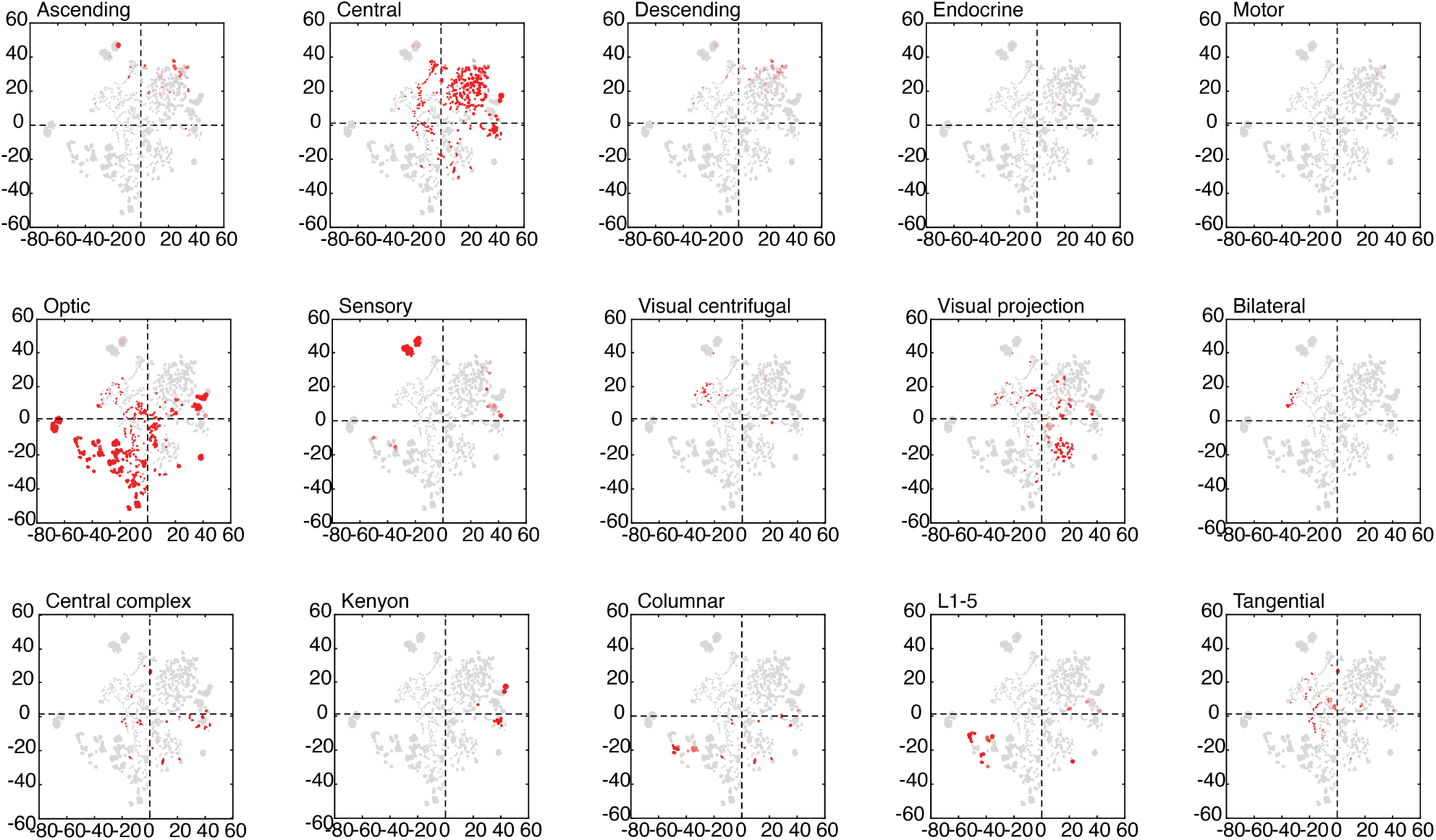
Communities in embedding space labeled by annotations. Here, we show communities in tSNE embedding space labeled by different annotations.

**Figure S8.**
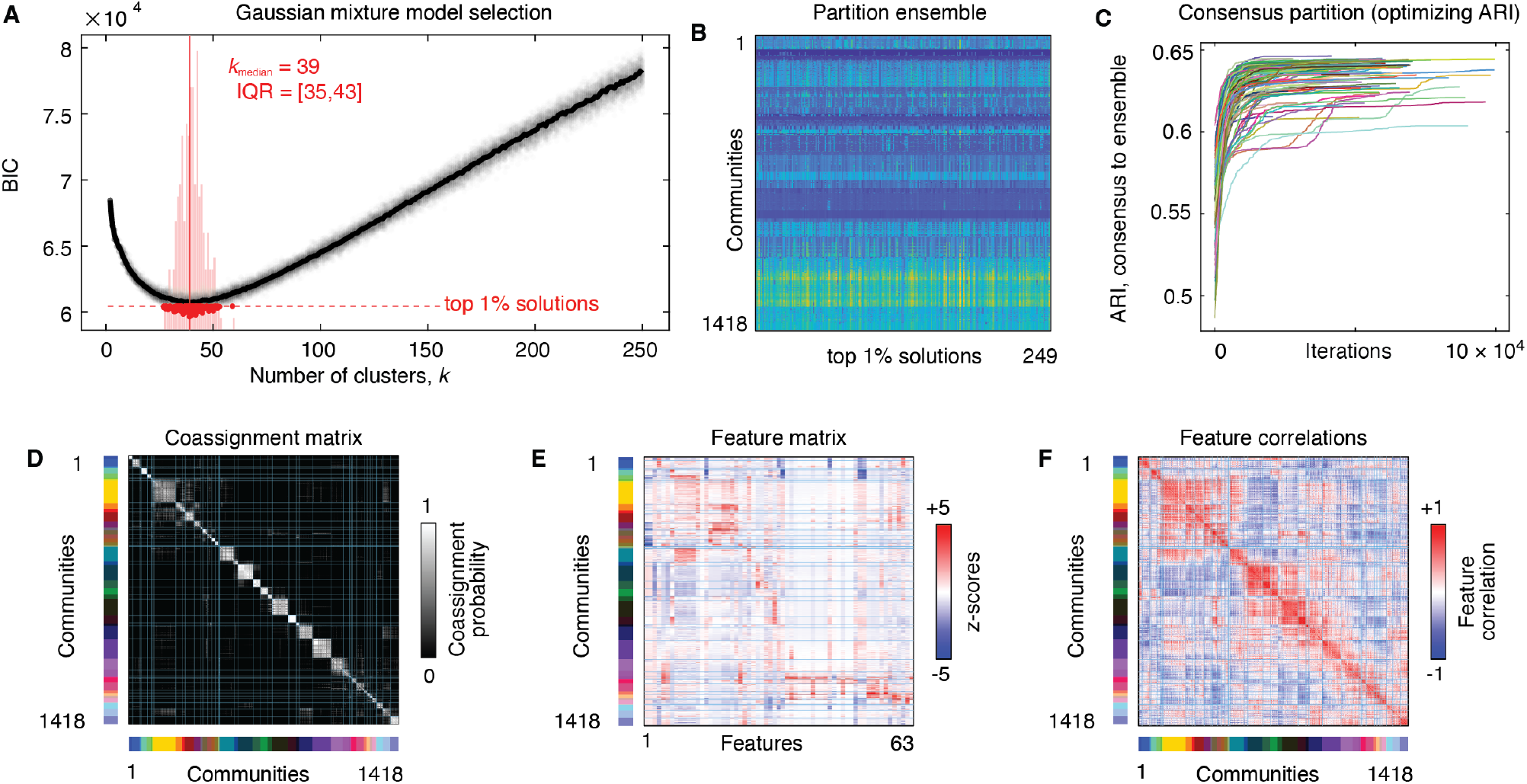
Fitting the GMM. We used a Gaussian mixture model to cluster communities based on their positions in PC space. Specifically, we varied the number of clusters from *k* = 2 to *k* = 250, repeating the optimization 100 times at each *k*. We ranked each solution in terms of BIC. (*a*) BIC as a function of *k*. We retained the top 1% of solutions based on BIC. The interquartile range was from *k* = 35 to *k* = 43 with a median of *k* = 39. (*b*) This procedure resulted in 249 partitions, which we refer to as the “partition ensemble.” In general, each of these solutions was good, in the sense that it represented an optimal balance of model complexity with model fitness. To obtain a point estimate or consensus partition, we used an optimization algorithm to identify a partition that was maximally similar (as measured by the adjusted Rand index; ARI), on average, to the partitions in the partition ensemble. We retained the partition with the maximum similarity. (*d*) Partition coassignment matrix ordered by consensus meta-clusters. (*e*) Community *×* feature matrix with rows ordered by consensus meta-cluster labels. (*f*) Feature correlation matrix with same ordering as the matrices shown in panels *d* and *e*.

**Figure S9.**
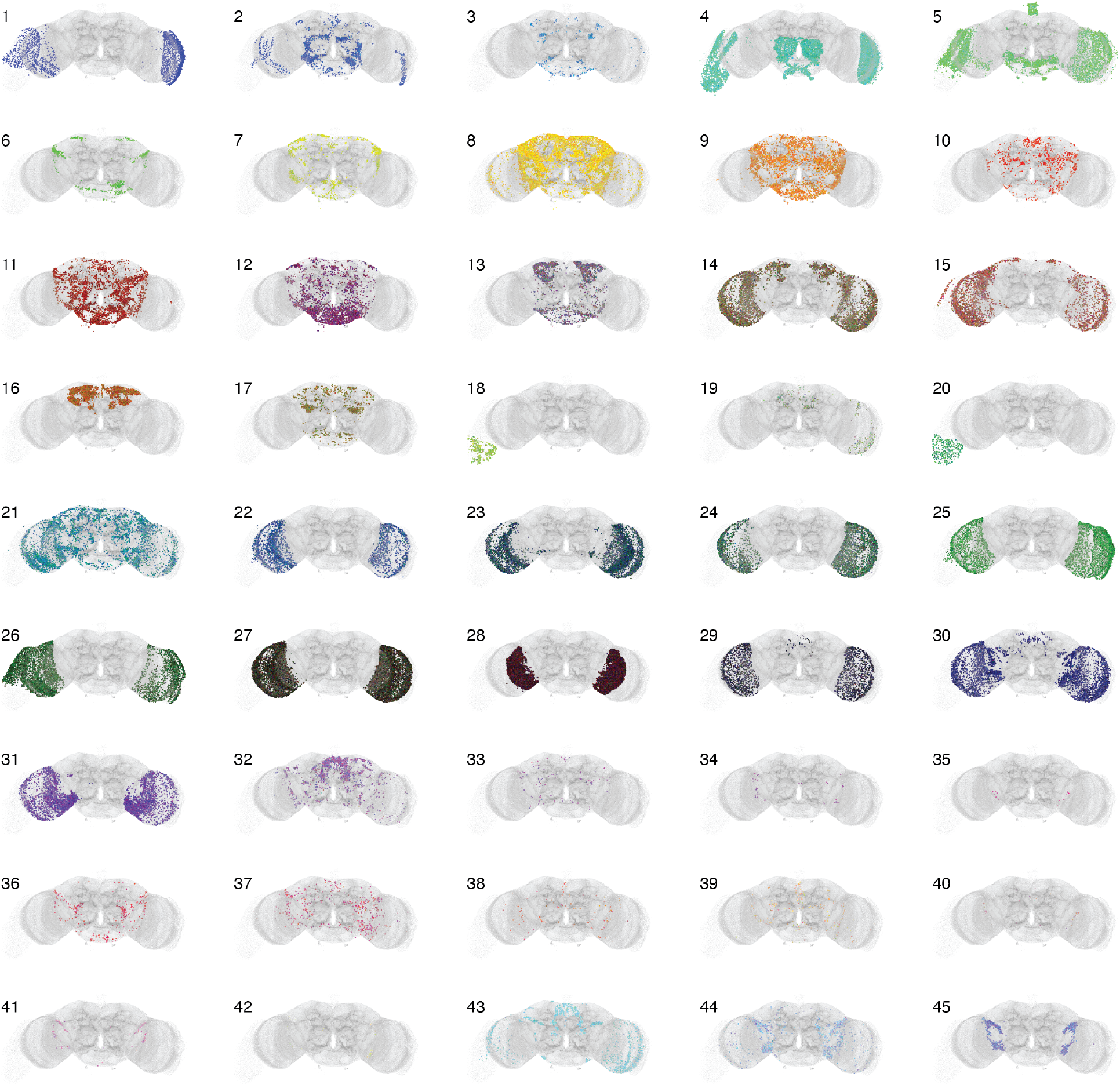
All 45 meta-clusters plotted in anatomical space.

**Figure S10.**
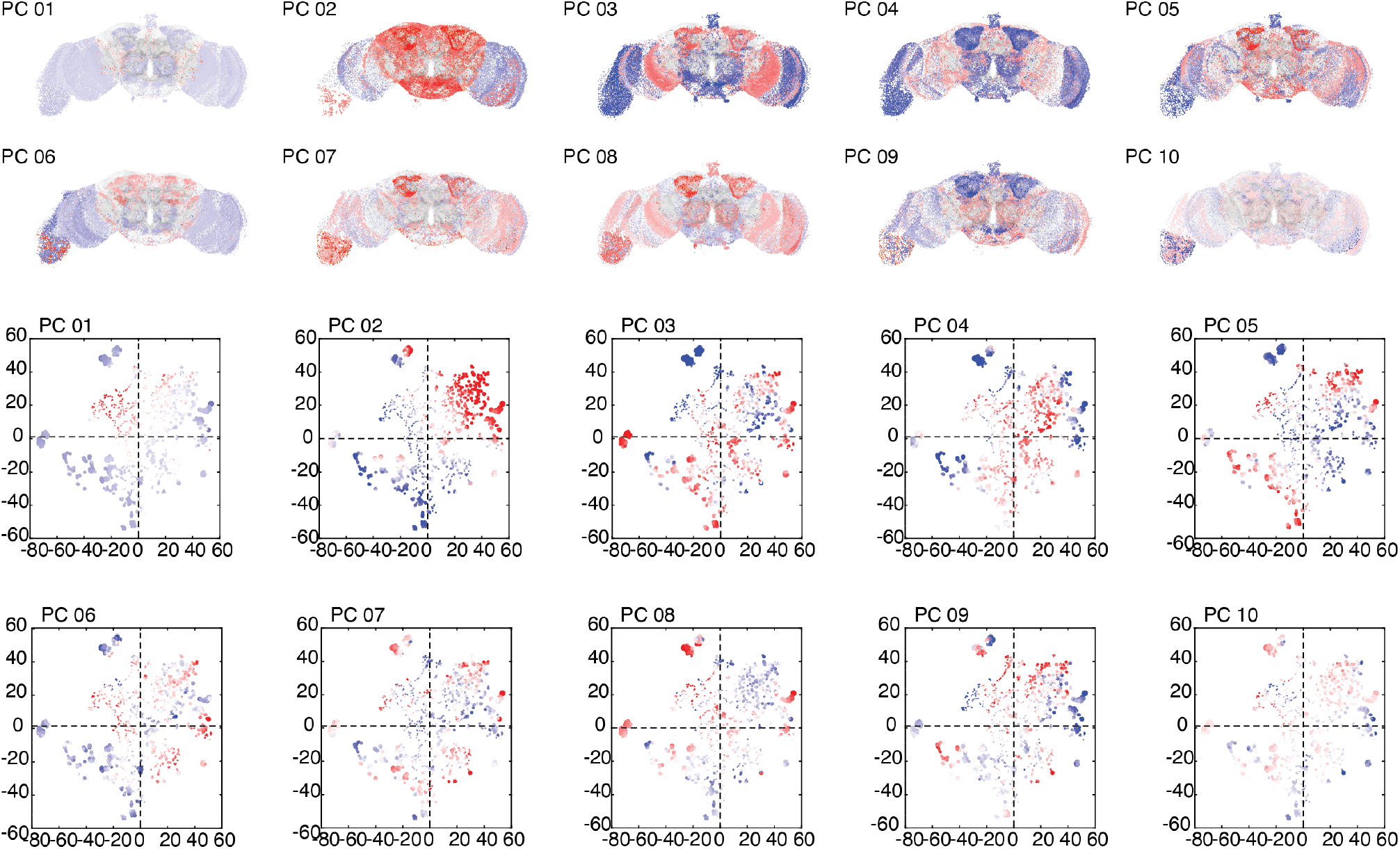
First ten principal components plotted in anatomical and embedding space.

**Figure S11.**
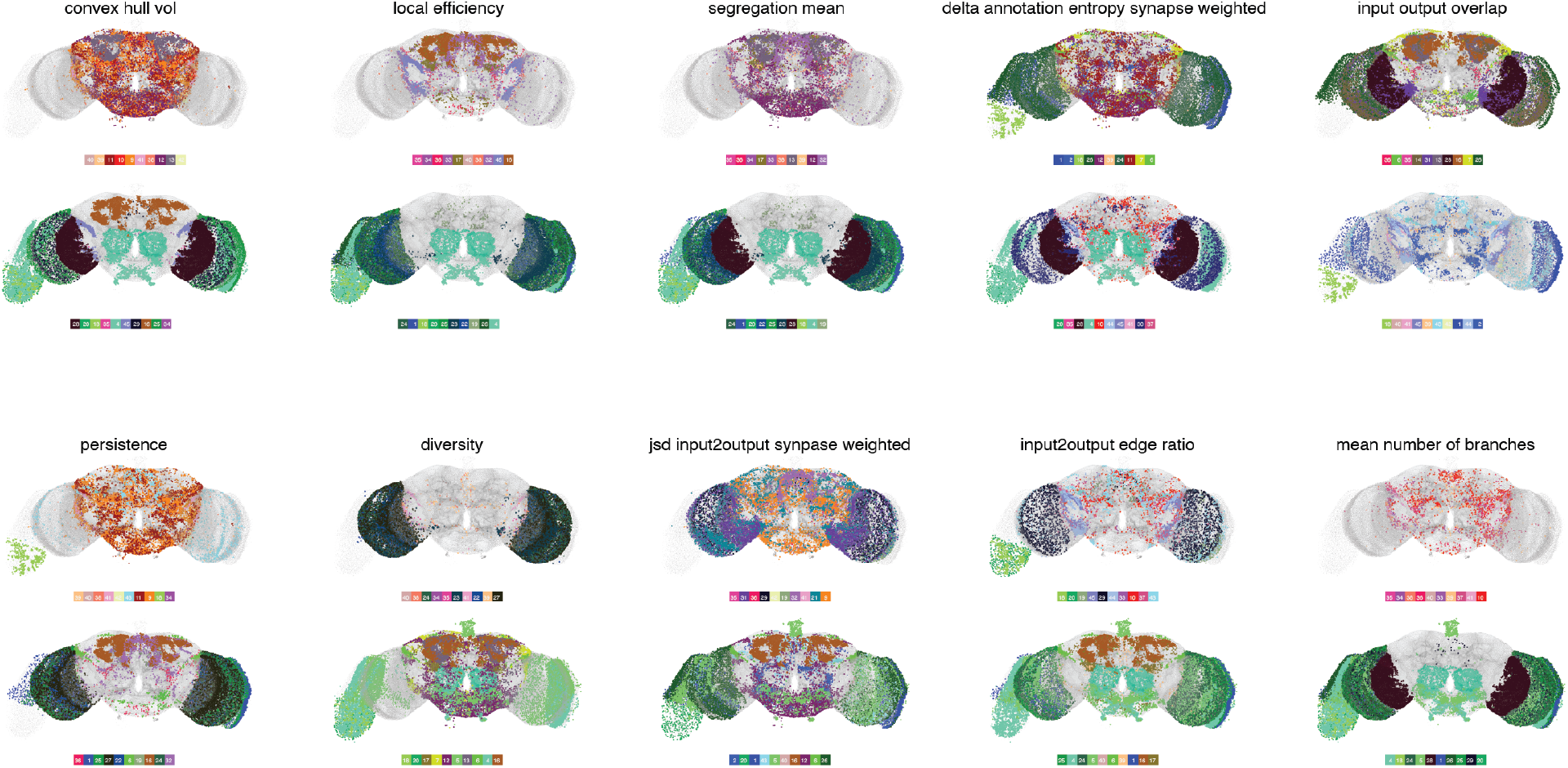
Extreme meta-clusters for a select set of brain features. For a select set of 10 features, we identified the top and bottom meta-clusters and plotted them in anatomical space.

**Figure S12.**
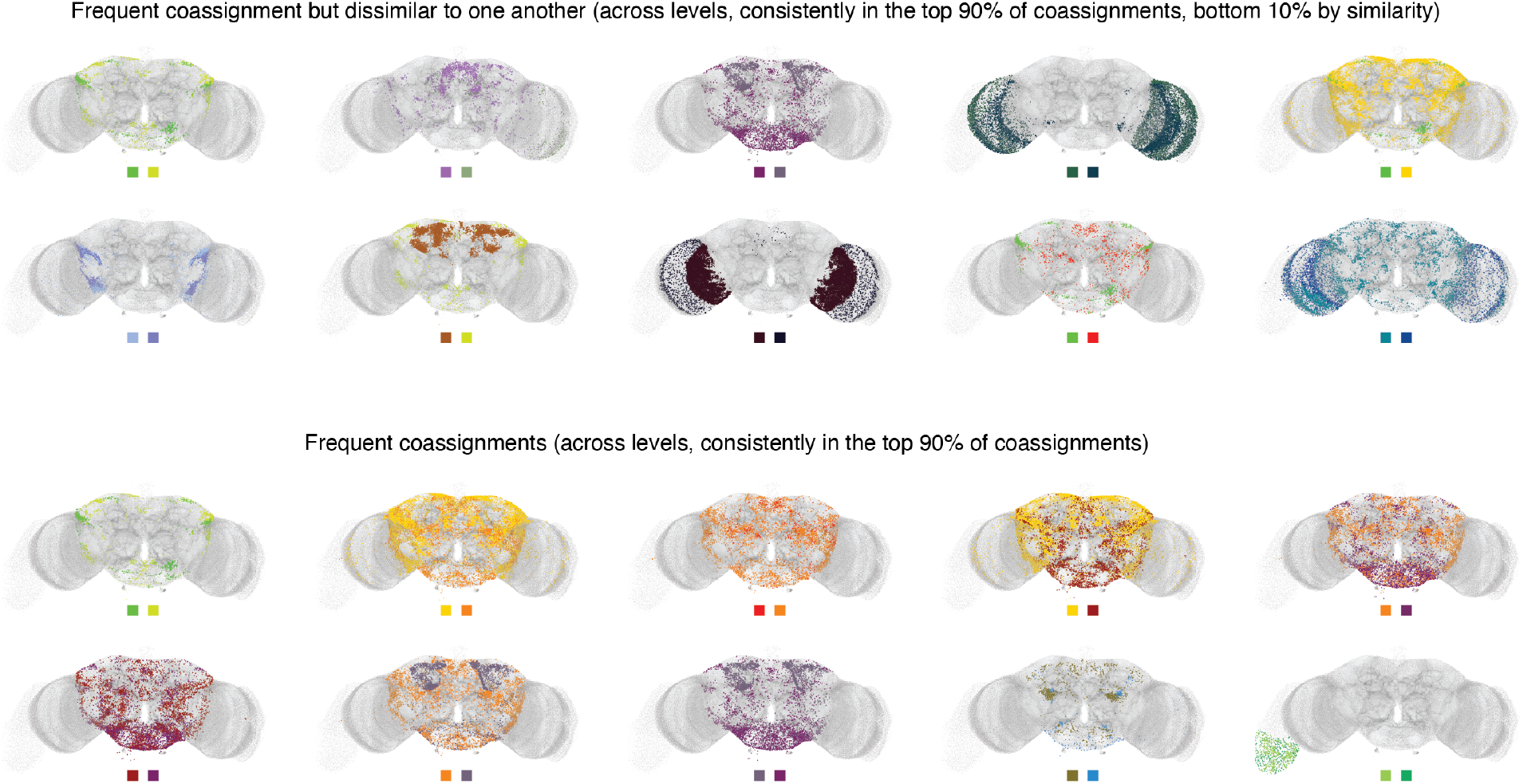
Frequently paired meta-clusters. We identified meta-clusters that were frequently coassigned to the same community across hierarchical levels. The top row shows meta-cluster pairs that were frequently coassigned to the same community (top 90%) but also exhibited dissimilar features (bottom 10%). The bottom row shows meta-cluster pairs that were frequently coassigned to the same community.

**Figure S13.**
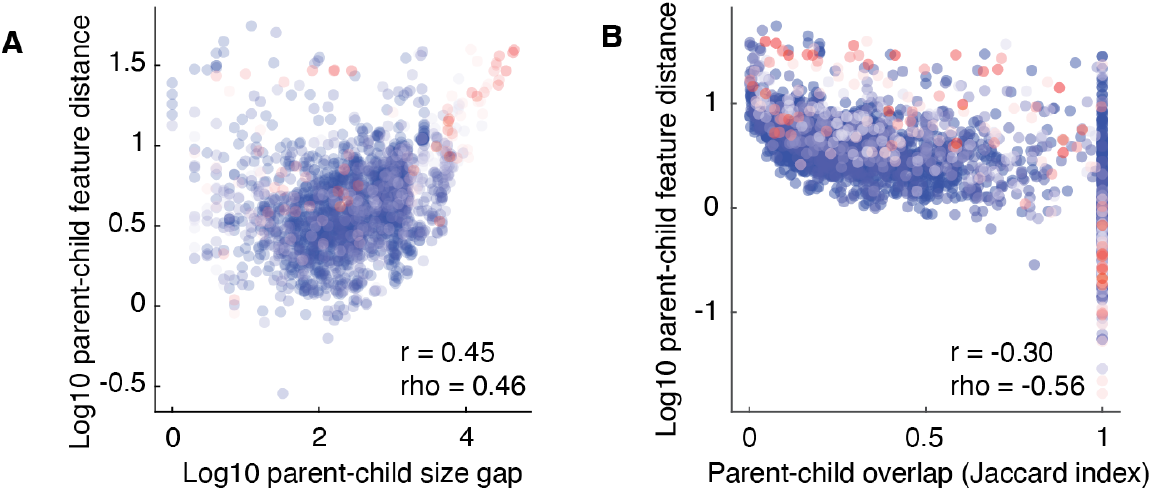
Linking parent-child feature distance to network statistics. We calculated the distance between parent and child communities in terms of their feature vectors. (*a*) We compared that value to the difference in size between parent and child communities. We found that they were positively correlated (*r* = 0.45, *ρ* = 0.46). (*b*) We also compared that value to the Jaccard index (overlap) between members of parent and child communities (*r* = −0.3, *ρ* = −0.56).

**Figure S14.**
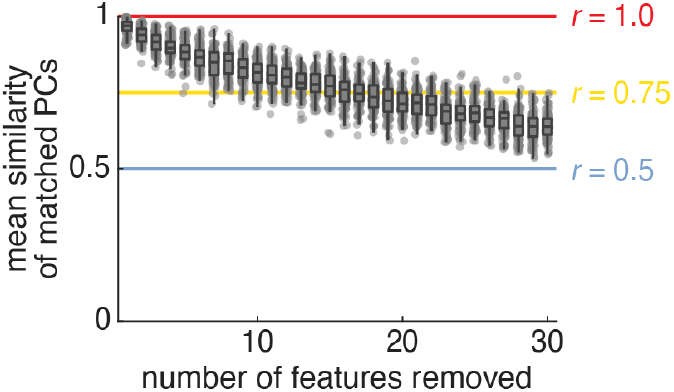
Meta-cluster sensitivity analysis. We assessed the impact of randomly removing different numbers of features on the PC landscape. After removing a subset of features, we calculated principal components using those that remained and aligned the resulting PCs to those reported in the main text using the Hungarian algorithm. We repeated this procedure for samples of size 1-30 (100 repetitions). This figure shows the mean alignment score (absolute correlation) of PCs as a function of the number of features removed. Each point represents an independent repetition.

